# The phase separation landscape of genome-wide genetic perturbations

**DOI:** 10.1101/2024.10.25.620319

**Authors:** Meta Heidenreich, Saurabh Mathur, Tong Shu, Ying Xie, David Sriker, Benjamin Dubreuil, Liam Holt, Emmanuel D. Levy

## Abstract

Biomolecular organization is central to cell function. While phase separation is a key mechanism orchestrating this organization, we lack a comprehensive view of genes that can globally influence this process *in vivo*. To identify such genes, we combined functional genomics and synthetic biology. We developed a bioorthogonal system that can identify changes in the intracellular milieu that globally tune phase separation. We measured *in vivo* phase diagrams of a synthetic system across >25 million cells in 2,888 yeast knockouts, and identified 68 genes whose deletion alters the phase boundaries of the synthetic system, an unexpected result given the system’s bioorthogonal design. Genes involved in TORC1 signaling and metabolism, particularly carbohydrate-, amino acid- and nucleotide synthesis were enriched. The mutants that changed phase separation also showed high pleiotropy, suggesting that phase separation interrelates with many aspects of biology.

**Highlights:**

- A synthetic protein system reveals the genetic and environmental tunability of protein phase separation
- Genetic knockouts affecting phase separation are highly pleiotropic
- Carbohydrate, amino acid, and nucleotide metabolism contribute to modulating phase separation potential
- Protein phase separation is a globally tunable property of the intracellular environment

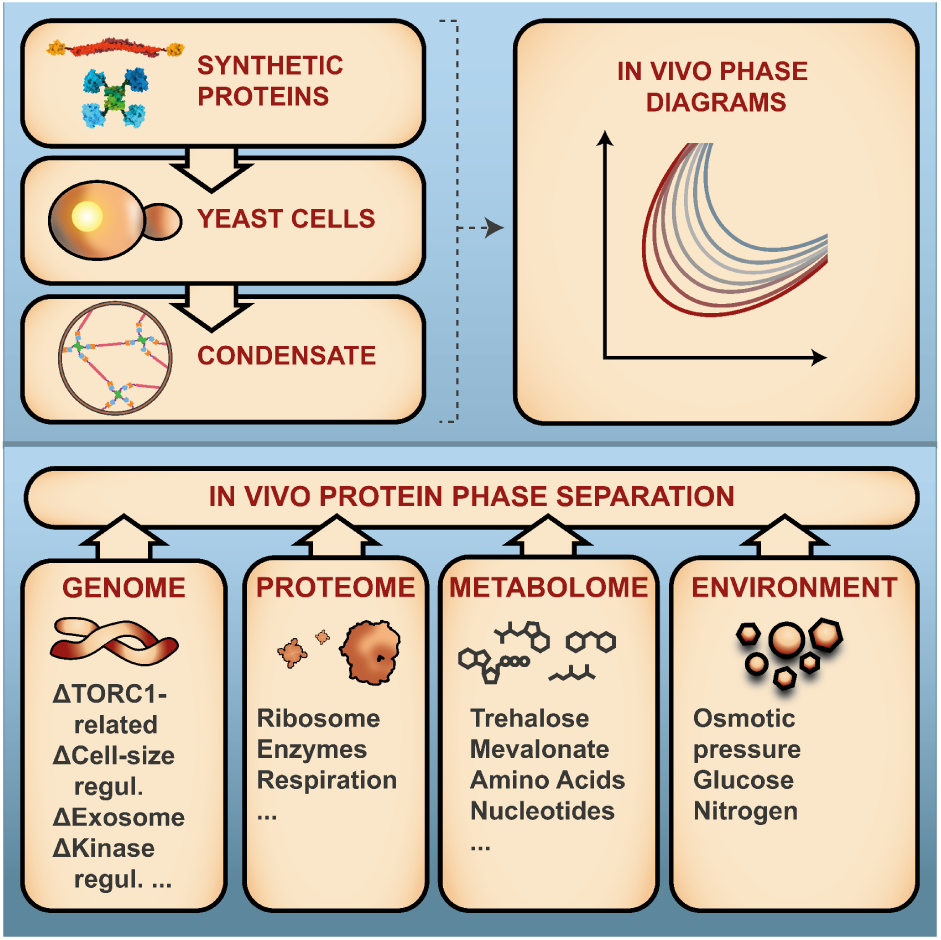

## Introduction

Phase separation has emerged as a fundamental mechanism of intracellular organization (*1–4*), and biomolecular condensation has been implicated in diseases, including neurodegenerative diseases, viral infections, and cancer (*5*, *6*). Therefore, identifying the intra-cellular factors that can influence the phase separation of proteins in cells is critical, both for fundamental research, e.g. to understand how the intracellular organization is orchestrated, and applied sciences, e.g., to decipher the mechanisms leading to diseases involving biomolecular condensates. For decades, researchers have studied how physical parameters, including temperature and pressure, impact phase separation. However, most of this physical and chemical knowledge is disconnected from biological systems where these parameters only vary to a limited extent. Nonetheless, growing evidence suggests that intracellular factors can globally modulate the phase separation behavior of biomolecules in cells (*7–10*). For example, the bacterial cytosol can undergo changes in material state depending on metabolic activity (*11*). Furthermore, studies increasingly show that the yeast cytoplasm undergoes extensive changes in its physicochemical properties depending on environmental conditions (*9*, *12–14*). For example, cells that are quiescent, energy depleted, or starved for carbon, show global rearrangements of their biomolecular content, and many proteins are observed to form large macromolecular assemblies under these conditions (*15–18*). However, it remains unclear whether these changes are driven by specific alterations in protein-protein interactions or if they arise due to more global physicochemical parameters such as pH (*19*), solvent quality (*20*), or molecular crowding (*21*). Additionally, the extent to which these physicochemical properties can be modulated by genetic alterations, such as gene knockouts, remains unexplored.

Here, we set out to investigate the genetic factors influencing protein phase separation using a synthetic system for which a phase diagram can be measured in living cells. We show that the system reacts as expected to osmotic compression. Furthermore, the system’s phase diagram changed dramatically in cells after entry into stationary phase, despite no changes in the apparent affinity between its constituent components. Due to the bioorthogonal nature of the system, these changes in phase behavior likely originated in a global regulation of intracellular protein phase separation propensity, independently of direct protein regulation. To map how genetic perturbations could impact such global regulation, we introduced this system to the yeast knockout (KO) collection and measured phase diagrams in thousands of single gene KOs, identifying 68 KOs that robustly change the system’s phase diagram. These KOs were enriched in TORC1-signaling and amino-acid homeostasis-related genes, and were associated with diverse phenotypes. In line with these findings, TORC1 inhibition by rapamycin, as well as nitrogen starvation influenced phase separation of our system in wild-type cells. Nitrogen starvation also inhibited phase separation of a different synthetic system that relies on interactions between disordered regions (*22*), and also decreased the tendency of the protein EDC3 to form condensates. These observations support that changes in the phase boundaries of our system reflect global changes in protein phase separation propensity, rather than specific effects on a particular constituent of the condensate, and that nitrogen availability and TORC1-signaling are important in maintaining the physicochemical properties of the cytoplasm. TORC1 signaling can influence protein phase separation by tuning ribosome-mediated crowding (*9*), however, diffusion measurement of 40-nm GEM particles showed that this mechanism does not explain the changes in phase separation in all KOs. Finally, metabolomic and proteomic profiling data of KO strains further highlighted the importance of the metabolism for the maintenance of phase separation homeostasis.

Taken together, our findings reflect an intricate interplay between genetic factors, cellular metabolism, and environmental cues in regulating intracellular protein phase separation.

## Results

### A synthetic minimal system can measure phase separation propensity using *in vivo* phase diagrams

We previously developed a synthetic system that allows its phase diagram to be measured in living cells. The system is composed of a tetramer and a dimer that interact with each other via binding domains to form mesh-like phase separated assemblies in living yeast cells (**Figure 1A-B**). Phase diagrams can be visualized for this system, by measuring the concentration of each component by proxy of green and red fluorescence intensities, and plotting the two concentrations against each other (*23*). If a condensate is present, dilute phase concentrations, i.e. concentrations outside the condensate, fall on the phase boundary and we refer to these as dimer-tetramer phase-boundary concentrations (PB-concentrations). By expressing the dimer and tetramer at stochastic levels, determined by random fluctuations in the copy number of separate expression plasmids, we can reveal their PB-concentrations across varying stoichiometries (**Figure 1C-D**). To test the responsiveness of our system, we artificially and rapidly increased the concentration of both components by reducing cell volume using hyperosmotic shock. After hyperosmotic shock, we observed widespread nucleation of the synthetic condensates, followed by progressive growth (**Figure 1E-I**, Movie S1 and S2). In this way, the percentage of cells with multiple puncta among those that had a condensate before the treatment increased over time, from 0.07% before the treatment to 0.42% after 15 minutes and 11.1% after 1h (**Figure 1G**). This delay indicates that newly nucleated condensates are not detectable shortly after the hyperosmotic shock due to their small size. We measured the phase diagram in cells before and after the hyperosmotic shock and observed an upward shift of the phase boundary 15 min after the treatment, as the dilute phase concentrations of dimer and tetramer increased. This shift was reversed one hour after the treatment, as condensates had matured and the dilute phase could be estimated reliably (**Figure 1H-I**). This result confirms that the system responds to concentration changes as expected from theory.

**Figure 1:**
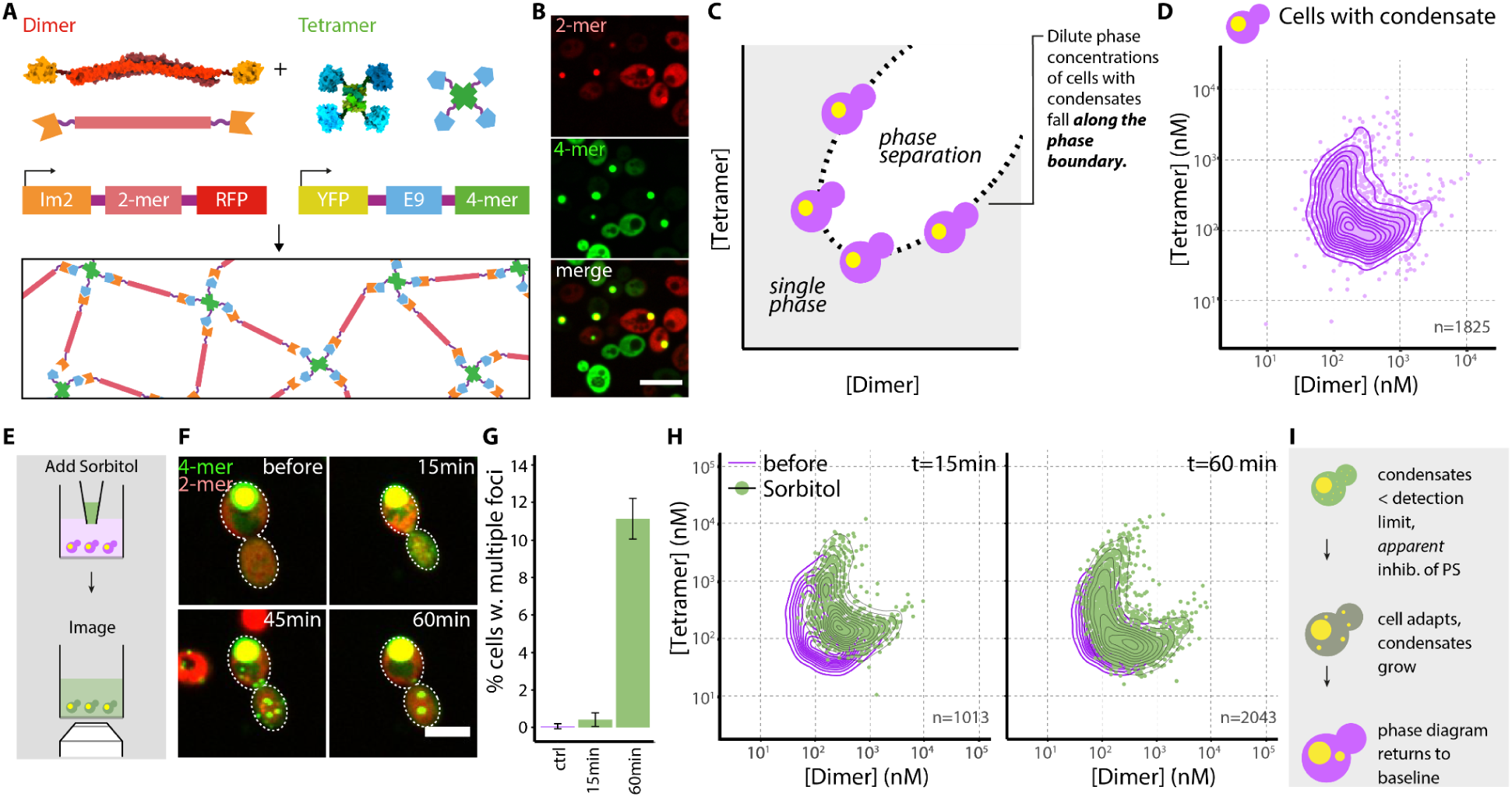
The previously developed synthetic protein system enables measuring changes in phase separation behavior. a. Schematic of the previously developed dimer-tetramer system. **A.** The system consists of two protein components, a dimer and a tetramer. Each component is encoded in a separate plasmid and carries an interaction domain (Im2 or E9), a multimerization domain (2-mer or 4-mer), and a fluorescent protein (RFP or YFP), respectively. A structural model of both components is shown along with a cartoon representation (fluorescent proteins are omitted for clarity). Schematics of the genetic constructs are shown underneath. The frame depicts the two components interacting and forming a mesh-like structure. **B. Representative images of cells expressing the two components at varying concentrations.** Red represents the dimer, and green the tetramer. Yellow highlights overlapping intensities. Scale bar=10μm. **C. Schematic of the phase diagram of our system**, as a function of the tetramer and dimer concentrations. Cells containing condensates will exhibit dilute-phase concentrations that are expected to fall on the phase boundary. **D. Phase diagram of the tetramer-dimer pair.** Each point represents the dimer- and tetramer concentrations of the dilute phase measured in cells with condensates; the solid lines show contours of the 2D density. **E. Schematic of the experimental setup for the hyperosmotic shock on cells expressing the system.** Cells in logarithmic growth phase were osmotically shocked with 1M sorbitol before imaging. **F. The cell volume decreases and condensates nucleate across the cytoplasm in response to an increase in concentrations of the system’s components.** Representative micrographs are shown prior to, and 15, 45 and 60 minutes after the hyperosmotic shock. Scale bar=5μm. **G. The fraction of cells with multiple condensates increases after hyperosmotic shock. H. A rapid increase in cytoplasmic concentrations of dimer and tetramer lead to a transient change in phase diagram.** Phase diagrams of cells immediately after the addition of 1M sorbitol (left) and 1h after the hyperosmotic shock (right) are shown. Green points represent tetramer and dimer concentrations of single cells, grey contour lines show the 2D density. Purple contour lines represent the 2D density of cells before addition of sorbitol. **I. A schematic of this interpretation.**

### Acute glucose starvation and stationary phase enhance phase separation

During glucose starvation, many proteins form punctate structures (*15*), and global physicochemical properties of the cytoplasm change, including a drop in pH (*24*), an initial fluidization (*25*) and then a transition to a solid-like state (*13*, *14*). Therefore, we next tested whether acute glucose starvation in wild type cells enhances phase separation of our system. Indeed, acute glucose starvation provoked a downward shift of the phase boundary (**Figure 2A**), and re-addition of glucose reversed this change with the phase boundary returning to its original position in the dimer-tetramer concentration space (**Figure 2B**). Previous works reported that global rearrangements of the cytoplasm during acute glucose starvation can be mediated by a decrease in cytoplasmic pH (*13*, *14*). Glucose starvation does not lead to intracellular acidification when the external pH is adjusted to 7.5 (*26*). However, acute glucose starvation in pH 7.5 media still shifted the phase boundaries of our system (**Figure 2C**), indicating that pH is one, but not the only parameter mediating the change of phase separation of our system during acute glucose starvation (**Figure 2D**). A similar effect was observed with cells in stationary phase where the dimer-tetramer PB-concentrations also underwent a downward shift (**Figure 2E**). Interestingly, in this condition, the range of concentrations at which phase separation took place widened greatly, suggesting larger cell-to-cell variability of phase boundaries compared to log-phase conditions (**Figure 2F**). Importantly, if the changes in phase separation are due to shifts in the intracellular milieu rather than chemical changes to the system components, the interaction between the components should remain the same. We previously showed that increasing the affinity between the dimer and tetramer resulted in a downward shift of the phase boundaries and longer fluorescence recovery after photobleaching (FRAP) profiles. By contrast, in stationary phase cells, the apparent affinity of our system did not significantly change as reflected in similar FRAP profiles. The dimer’s relative recovery after 25 s was 0.66±0.02 during growth and 0.69±0.05 in stationary phase, while the tetramer’s recovery was 0.64±0.03 and 0.62±0.05, respectively (**Figure 2G-H**). Thus, the shift in the phase boundary is likely due to global changes in the physicochemical properties of the cytoplasm rather than specific changes in protein interaction.

**Figure 2:**
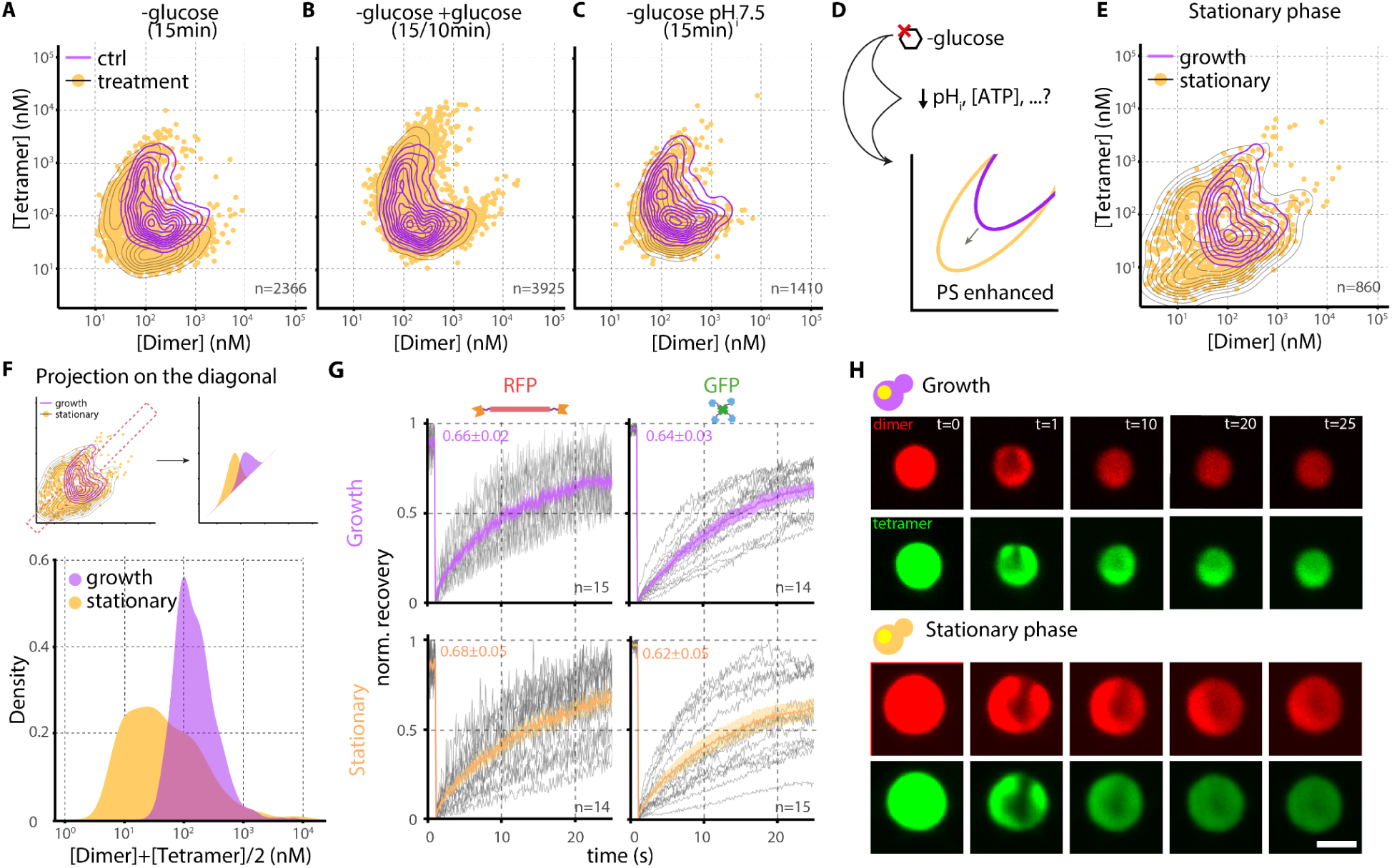
The phase separation behavior of our synthetic system changes during acute glucose starvation and in stationary phase. **A-B. Acute glucose starvation enhances phase separation reversibly.** The phase diagram of cells that were starved for glucose for 15 min (A) shows a downward shift of the phase boundary, while re-addition of glucose reverses this effect (B). Orange points represent the dimer-tetramer PB-concentrations after treatment, grey contour lines show the 2D density, and purple contour lines show the 2D density of control cells washed with media containing glucose. The same control data is shown in A and B (n=1138). The number of cells in each treatment is indicated in the inset. **C. Glucose starvation without acidification also enhances phase separation.** Cells were washed into pH7.5 media with (purple) or without glucose (orange) and imaged after 15 min. **D. Schematic highlighting that glucose starvation enhances phase separation partly via intracellular acidification. E. Phase separation is enhanced in stationary phase.** Dimer and tetramer PB-concentrations of cells grown to stationary phase for three days (orange points, grey contour lines) or in logarithmic phase (purple contour lines).The number of cells is indicated in the inset. **F. Stationary phase leads to an increased range of concentrations at which phase separation occurs, implying cell-to-cell heterogeneity of phase boundaries.** The data from panel E was projected onto the diagonal, highlighting that PB-concentrations of cells in logarithmic phase (purple) span a narrower range than those in stationary phase (orange). **G-H. The diffusion of dimer (RFP, left) and tetramer (GFP, right) within condensates remains unchanged in cells reaching stationary phase. G.** Results of FRAP experiments on condensates in logarithmic (upper panel, purple) or stationary phase cells (lower panel, yellow). The recovery in the RFP channel (dimer, left column) and GFP channel (tetramer, right column) are shown. Grey lines represent measurements of individual condensates, solid lines indicate the mean of each experiment, transparent areas indicate the standard errors. Mean and standard error after 25 seconds of recovery are indicated. **H.** Representative images of condensates during FRAP experiments. Scale bar: 2μm.

These results highlight that acute glucose starvation enhances phase separation, with pH contributing, but not solely accounting for this effect. Additionally, cells in stationary phase display a wider range of phase separation propensities than cells in log-phase, possibly reflecting a bet-hedging strategy in the physicochemical state of the cytoplasm.

### A yeast library to measure the genetic factors influencing phase separation

To investigate genetic factors influencing protein phase separation propensity, we introduced the synthetic system to the yeast knockout (KO) library by synthetic genetic array (SGA) (*27–29*). In the resulting library, each strain expresses the synthetic system in the background of a specific single gene deletion (**Figure 3A**). We imaged the resulting library, composed of 3022 strains, acquiring 193,536 images, and quantifying the concentrations of the dimer and tetramer in >25 million segmented cells. To analyze thousands of phase diagrams, we employed two methods. For the first method, we fitted an ellipse to the PB-concentrations of a strain and used the ellipse center to summarize its location (**Figure 3B-C**). The resulting density distribution of ellipse centers peaked at a particular coordinate. We defined strains close to this peak density as “reference strains”, whereas strains farthest from this peak density were identified as potential outliers (**Figure 3D**). This “center-method” measures only the displacement of the entire phase boundary and we therefore employed a second analysis method that could capture changes in the scaling of the ellipse, as such changes would not necessarily change its center. This method considers the upper and lower parts of the ellipse as two separate “branches” and we refer to this second method as the “branch-method” (Methods, Supplementary Figure 1). We collected the strains exhibiting the most extreme changes PB-concentrations based on either method, as well as reference strains from the center of both distributions. This set of 301 KOs comprises 186 identified by the ellipse center-method and 189 by the “branch-method”, of which 74 were common. The outlier and reference strains were transferred to a new 384-well plate, which was imaged and analyzed three more times. Both measurements showed good correlations across the repeats (Pearson correlation coefficients between the first and second repeat: 0.89, and 0.90 for x and y coordinates of the ellipse fit’s centers, and 0.83 and 0.87 for the second method, Supplementary Figure 2). The 68 strains that behaved as outliers in at least two repeats were deemed high confidence outliers (**Figure 3E**, Supplementary Figure 3). Interestingly, among those KOs, five are encoded on the complementary DNA strand of another high confidence outlier (YPT6/VPS63, SLM6/REI1, YNL198C/GCR2, YML012C-A/UBX2, YKR047W/NAP1). As there are 303 overlapping ORFs in the library, we expect to sample less than one by chance from a draw of 68, so observing five represents a significant enrichment (Wilcoxon test *P*<2.2*10^-16^).

**Figure 3:**
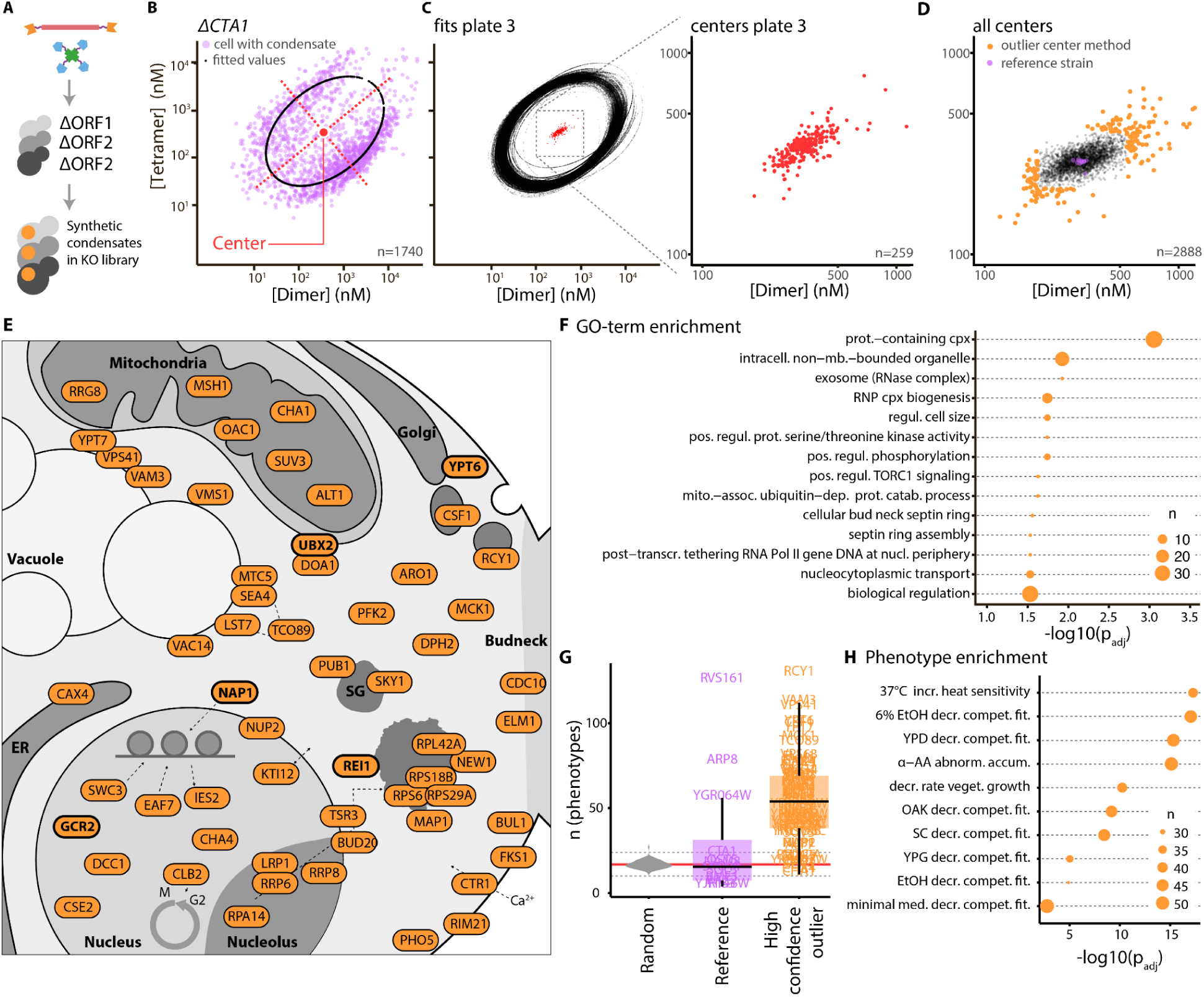
A genome wide screen to identify KO strains with changed phase diagrams of our system A. Schematic of the experimental setup. The synthetic system was introduced into the yeast knock out (yKO) library, enabling the measurements of phase diagrams in the background of thousands of KOs. **B-C. Analysis of phase diagrams using the hysteresis package in R**. First, an ellipse is fitted to the phase diagram of each strain. The fit for the reference strain ΔCTA1 is shown (B). Purple points represent cells with a condensate, black dots represent the elliptic fit. The red point highlights the center of the ellipse fit. All fits for one plate of the library are depicted in C. The red points in the center indicate the center points of the ellipses. A magnification of the plot is shown on the right. **D. Result of the ellipse fit-based analysis of the screen.** Center points of fitted ellipses are shown for all knock out strains. From this method, 186 outlier strains (orange) and eleven reference strains (purple) were selected for further validation. **E. Schematic overview on genes that change phase separation.** Bold genes have an overlapping ORF among the high confidence outliers. 58 of 68 ORFs are illustrated. **F. GO term analysis** of KOs leading to changed phase separation (n=68). The adjusted *P*-value for the enriched GO terms is shown, the size of the dot indicates the number of genes associated with each term. The analysis was performed with gProfiler (*36*), the *P*-value was adjusted via Benjamini-Hochberg correction. prot. = protein, cpx=complex, intracell.=intracellular, mb.=membrane, regul.=regulation, pos.=positive, mito.=mitochondria, assoc=associated, dep.=dependent, catab.=catabolic, transc.=transcriptional, nucl. = nuclear. **G. KOs identified as influencers of phase separation are highly pleiotropic**. The median number of phenotypes was calculated for a random set of 68 KOs of the strains screened. This was repeated 100 times and the distribution of medians is shown as a violin plot (gray). The mean of this distribution is shown as a solid red line across the plot. The dotted lines highlight the mean ± 3* standard deviation. The set of KO strains from the center of the distributions of our screen’s results (purple) is not significantly different from this distribution (purple, Welch’s t-test *P*=.30), while the 68 outliers show significantly more phenotypes (Welch’s t-test *P*<2.2*10^-16^). Boxes delineate the first and third quartiles, the middle line corresponds to the median, the upper and lower whiskers extend to the highest and lowest values, at most 1.5 times the interquartile range. **H. Phenotypes enriched in KOs that change phase separation of our system.** The phenotypes associated with a random set of 68 KOs were sampled 100 times and the observed and randomly expected frequencies of each phenotype were compared. The size of the dot indicates the number of outlier KOs annotated with each phenotype. Significance was tested using Z-test with Benjamini-Hochberg *P*-value adjustment and the -log10 of the adjusted *P*-value is shown. Incr. = increased, decr. = decreased, compet. = competetive, fit. = fitness, AA = amino acid, abnorm. = abnormal, accum. = accumulation, veget. = vegetative, med. = media.

Furthermore, 10 dubious ORFs were detected as high confidence outliers, however, only YLR402W did not have an open reading frame on the complementary strand or immediately next to it. YLR402W does not have any ortholog, even in closely related Saccharomyces species, but it is present in a dataset of 1,000 *S. cerevisiae* strains (*30*). We find that expressing this protein rescues the change in phase boundary observed in the KO. Interestingly, structure prediction suggests YLR402W can self-assemble into amyloid-like structures and its expression modulates cell size (Supplementary Figure 4). Altogether, these data imply that YLR402W is a functional protein and we propose to call it AIS1 for “Altered Intracellular phase Separation”.

All high confidence outliers and their details are listed in Supplementary Table 1. Gene Ontology (GO) term analysis (*31*, *32*) showed that “protein complex” was a term over-represented among the high confidence outliers along with “membrane-less organelles”, “cell size regulation”, “regulation of phosphorylation”, and “TORC1-signaling”, among others (**Figure 3F**).

We next examined the phenotypes associated with the KOs as annotated in YeastMine (*33*). Outlier strains are highly pleiotropic and are associated with a disproportionately large number of phenotypes (median=54) relative to random sets of KO strains (median=16-17; 16.23 on average across 100 samples, two-sided Z-test *P*< 2.2*10^-16^, **Figure 3G**). Similarly to the random sets, reference KOs that did not exhibit a change in PB-concentrations and showed equally few annotated phenotypes (median 15.5, two-sided Z-test *P*=.80, **Figure 3G**). Interestingly, among the enriched phenotypes, *“abnormal alpha-amino acid accumulation”* was associated with 50 outliers of our screen (74% of total), while this phenotype was only represented in 29% of strains in a random set. Similarly, *“increased heat sensitivity”* was associated with 38 outliers (56% of total) compared to 17% in the random set (**Figure 3H**, Supplementary Table 2).

Finally, we tested the impact of the high confidence outliers on the formation of mesoscale assemblies by tagging and imaging EDC3 (P-bodies (*34*)) and HSP104 (JUNQ/IPOD (*35*)) in these KO strains. When compared to reference KO strains, the outliers showed a significantly higher fraction of cells with puncta of endogenously tagged EDC3 (*P*=.0046), while the fraction of cells with puncta of HSP104 was not significantly altered (*P*=.1297). However, the variance of the fraction of cells with puncta was significantly altered in both cases (F-test *P*=.0008 (EDC3) and *P*=.0010 (HSP104), Supplementary Figure 5). Thus, the KOs modulating the phase separation of the synthetic system also show a disproportionate impact on the propensity of EDC3 and HSP104 to self-assemble at the mesoscale.

### TORC1 inhibition, nitrogen and amino acid starvation globally inhibit protein phase separation

The GO-term ‘*positive regulation of TORC1 signaling*’, and the phenotype *“abnormal alpha-amino acid accumulation”,* to which TORC1 signaling is responsive, as well as the phenotype *“decreased resistance to [the TORC1 inhibitor] sirolimus”* were enriched in the high confidence outliers of our screen. Furthermore, the protein complexes represented with multiple subunits in the high confidence outliers could be linked to TORC1 signaling (**Figure 4A**). This prompted us to test whether TORC1 inhibition would change the phase diagram of our system in wild type cells. Indeed, TORC1 inhibition with 100 nM rapamycin inhibited phase separation, whereby higher concentrations of both components were necessary for phase separation after four and six hours, but not after two hours of treatment (**Figure 4B**). TORC1 signaling is responsive to nitrogen availability, and cells starved for nitrogen for six hours also showed inhibited phase separation of our system, as seen by an upward shift of the phase boundary in the phase diagram (**Figure 4C**). To test whether this inhibition of phase separation is a general phenomenon, we conducted the same experiment with two additional phase separating systems: a second synthetic system based on intrinsically disordered regions interactions (*22*) (**Figure 4D-F**), as well as the naturally phase separating protein EDC3 (**Figure 4G-H**). Both were expressed from plasmids to yield a high variability in expression, and both systems showed inhibited phase separation after six hours of nitrogen starvation, as seen by an increased dilute phase concentration (**Figure 4D-H**).

**Figure 4:**
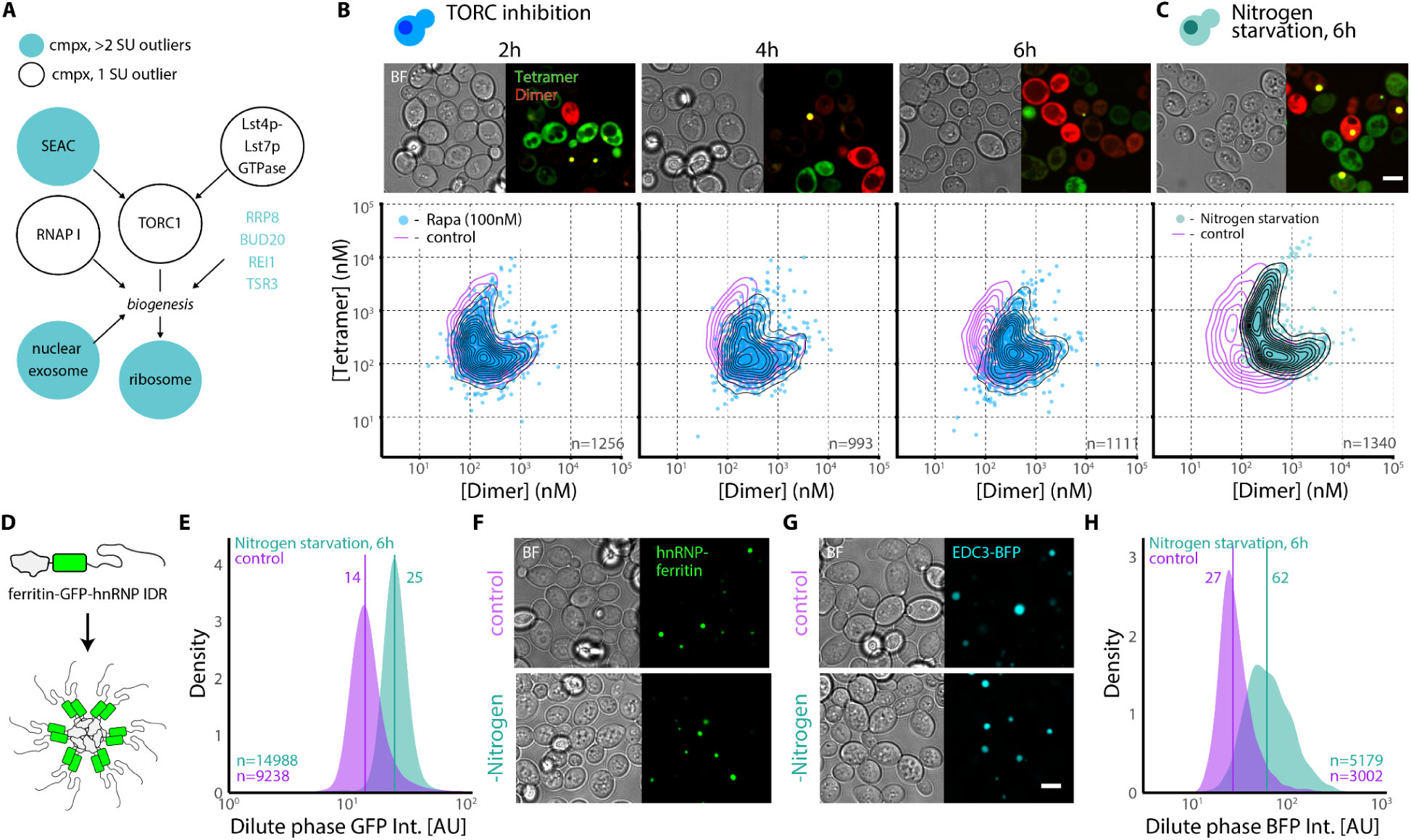
TORC1 activity influences *in vivo* phase separation. a. Schematic showing how TORC1 signaling and ribosome biogenesis are connected, with respect to our outliers. **A.** The four complexes that are represented with two or more high confidence outliers (ribosome, SEACAT, SWR1-c, and nuclear RNA exosome, turquoise circles) all play a role in TORC1 signaling-mediated ribosome biogenesis. Other TORC1-signaling-related genes that are high confidence outliers in our screen are also shown (white circles). Arrows indicate indirect or direct relationships. cmpx= complex, SU = subunit. **B. TORC1 inhibition changes phase separation behavior of the system in a time-dependent manner.** Growing cells were treated with 100 nM rapamycin and imaged after two, four and six hours (from left to right). The phase diagram of treated cells (blue) is compared to that of control cells treated with DMSO (purple 2d-density) **C. Nitrogen starvation decreases phase separation of our system.** The phase diagram measured in cells after 6h nitrogen starvation (turquoise points, dotted line represents the 2D density) is compared to that seen in log-phase cells (purple, 2D density). A representative micrograph of cells in nitrogen starvation is shown above. Scale bar= 5 μm. **D-F. An orthogonal synthetic protein system also shows decreased phase separation in nitrogen-starved cells. D. Schematic of the orthogonal system** (*22*). The orthogonal one-component system consists of a ferritin-derived dodecamerization domain, GFP and an intrinsically disordered interaction domain from hnRNP. **E. Nitrogen starved cells require a higher minimum concentration for phase separation of the orthogonal system.** The density of the low-density phase GFP intensities of the system in growing cells (purple) and cells in nitrogen starvation (6h, turquoise) is shown. The median intensity of the two conditions are highlighted by solid vertical lines, the median values are indicated (Welch two-sample t-Test *P*<2.2*10^-16^). **F. Representative micrographs of cells expressing this system during log phase and nitrogen starvation**. **G-H. The naturally phase separating protein EDC3 also shows decreased phase separation in nitrogen starvation.** Cells expressing EDC3-BFP from a plasmid were subjected to nitrogen starvation. Representative micrographs are shown in H. Scale bar=5μm. The density of BFP intensity in the dilute phase of cells with an EDC3 puncta in nitrogen starvation (turquoise), and during logarithmic phase (purple) are shown. The median intensity of the two conditions are highlighted by solid vertical lines, the median values are indicated (Welch two-sample t-Test *P*<2.2*10^-16^).

### The change in phase separation of selected KOs is likely influenced by crowding and metabolism

TORC1 has been shown to alter phase separation by controlling crowding through changes in ribosome density (*9*). To investigate whether ribosome-mediated crowding underlies the changes in phase boundaries in our system, we selected a set of six KOs that altered the system’s phase behavior (ΔNAP1, ΔLST7, ΔUBX2, ΔYML010W-A, ΔVPS41 and ΔYLR402W), and assessed their intracellular crowding using 40 nm genetically encoded multimeric nanoparticle (GEM) diffusion measurements (**Figure 5A**). ΔNAP1, ΔUBX2, ΔYML010W-A and ΔVPS41 were chosen because they influenced the phase behavior of our system strongly, ΔLST7 because it is a known positive regulator of TORC1, and ΔYLR402W because it is the only ORF of unknown function with no other gene encoded on the opposite strand in the genome. In these measurements, GEM nanoparticles are tracked inside the cells, and their diffusivities are used to infer intracellular crowding on the 40 nm length scale. GEM diffusivities were significantly increased in ΔLST7 and ΔΥLR402W compared to the reference strain ΔHIS3 (ΔLST7: 0.25±0.01 μm^2^ s^−1^, ΔΥLR402W: 0.26±0.02 μm^2^ s^−1^, ΔHIS3: 0.17±0.01 μm^2^ s^−1^), while they were slightly decreased in ΔNAP1 (0.16±0.01 μm^2^ s^−1^), and not significantly altered in the other KO strains (**Figure 5B**). We then measured the diffusivities of our condensates to explore whether changes in phase separation propensity were associated with differences in crowding at the mesoscale. Five of the six strains indeed showed altered motion of the condensates. Specifically, ΔNAP1, ΔLST7, and ΔYLR402W showed increased diffusivities, while ΔUBX2 and ΔVPS41 showed the opposite effect (**Figure 5C**).

**Figure 5:**
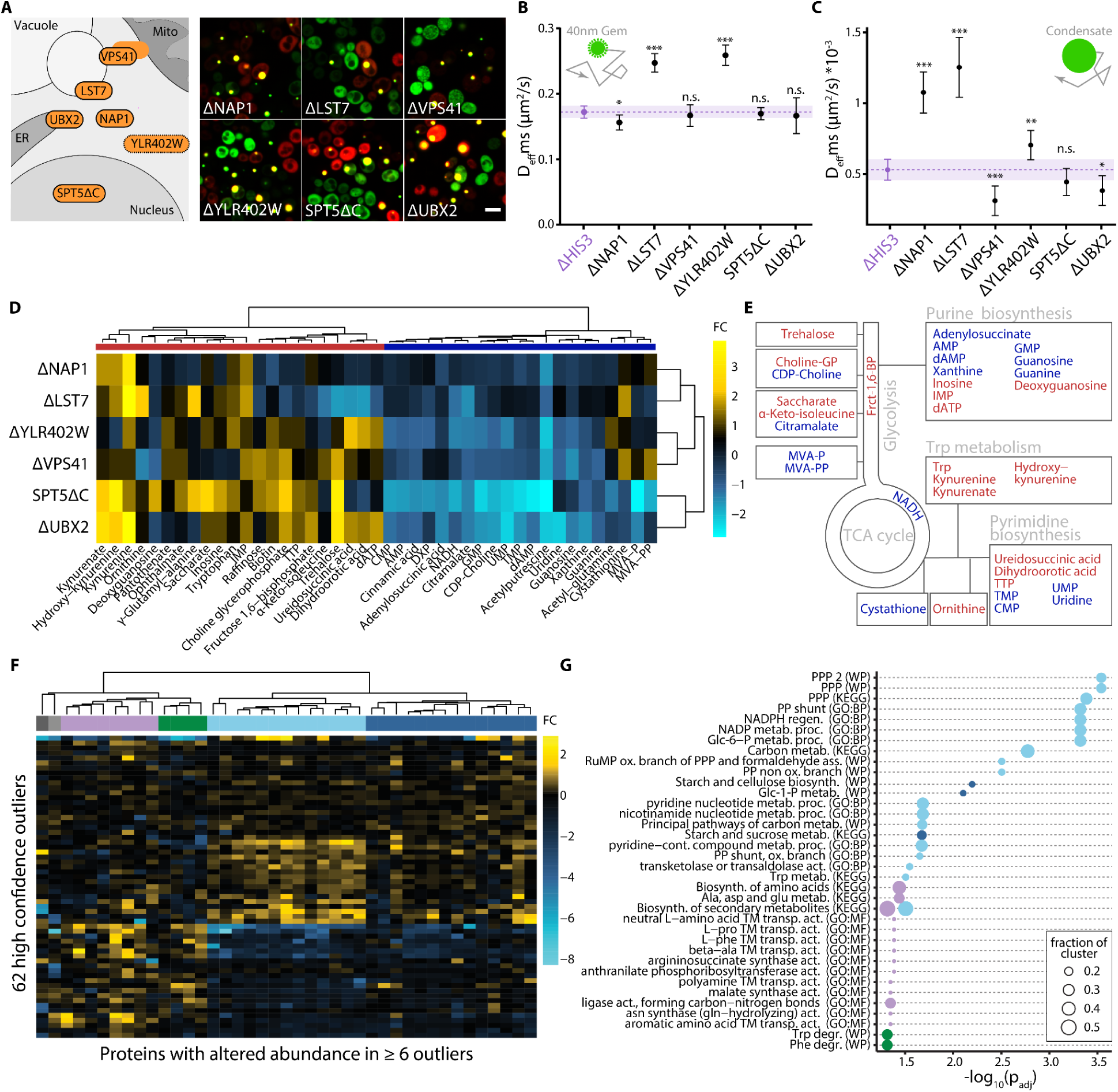
An in depth analysis of mutants that impact phase separation reveals complex effects on cytoplasmic dynamics, proteome abundance, and metabolism. A. Schematic overview of six selected outliers from our screen and representative images of the selected KOs with our synthetic system. Scale bar = 5 μm**. B. Nanorheology measurements using 40nm-GEMs for these six KO strains.** Average diffusivity of 40nm-GEMs per cell was measured and compared to the diffusion in ΔHIS3 (purple). Per cell, the diffusion of at least three tracked GEM nanoparticles was averaged. Points show the mean diffusion, error bars indicate the 95% confidence interval. The dotted line highlights the mean diffusion in the reference strain ΔHIS3, the purple shaded area indicates the 95% confidence interval of the diffusion in this strain. Welch’s two-sample t-test: n.s.: not significant, * *P*<.05, \*\*\**P*<.001, n.s. *P*>.05**. C. Diffusivity of condensates.** The diffusivities of the condensates were measured and compared to ΔHIS3 (purple). * *P*<.05, \*\**P*<.01, \*\*\**P*<.001, n.s. *P*>.05. **D. Metabolomic profiling of the six strains.** The heatmap shows fold changes of metabolites significantly changed in at least two strains, compared to ΔHIS3. **E. Schematic of the metabolites and associated pathways detected in D. F.-G. Analysis of proteomics data reported in** (*38*)**. F. Heatmap of protein abundance (x-axis) for 62 outliers (y-axis) present in the dataset.** The proteins shown are those differentially expressed (>2-fold) in at least six outlier strains relative to reference strains. The heatmap highlights four clusters and two singletons (gray). **G. GO-term enrichment analysis of the clusters shown in F.** The analysis was performed on each cluster, using gProfiler (*36*), Benjamini-Hochberg correction was used for *P*-value adjustment. Only GO-terms associated with at least 10% of the proteins in the cluster are shown. The size represents the fractions of proteins in each cluster associated with the respective GO-term, colors indicate the corresponding cluster shown in F.

To further characterize this subset of KOs and identify potential intracellular effectors of phase separation propensity, we profiled the metabolome of these six strains. Out of 166 metabolites detected in all samples, the abundance of 43 compounds was significantly different from the control strain ΔHIS3 in at least two strains. These were mostly related to the nucleotide-, tryptophan-, phospholipid-, and mevalonate-metabolism, as well as notably trehalose, a sugar previously shown to alter viscosity of the cytoplasm (*37*) (**Figure 5D-E**). The importance of the metabolism was further reflected in a dataset of differentially expressed proteins reported in (*38*). Among the 68 KOs that changed phase separation, 62 were profiled in this study. To characterize commonalities in their proteomes, we first identified proteins disproportionately changing abundance among them. We identified 41 such proteins out of 1850 (Methods), and these mapped to four clusters (**Figure 5F**) enriched in the terms “*Pentose-phosphate pathway*” (Wikipathways (WP) (*39*)), “*Starch and cellulose biosynthesis*” (GO), “*Biosynthesis of amino acids*” (KEGG (*40*)), and “*Tryptophan/Phenylalanine degradation*” (WP), respectively (**Figure 5G**). Finally, our own proteomics profiling of the six strains used for metabolics also highlighted the importance of the metabolism (**Figure S7**).

## Discussion

Our synthetic two-component system enabled high-throughput characterization of *in vivo* phase diagrams and reacted to intracellular concentration changes as expected: nucleation of droplets increased after osmotic compression of cells (**Figure 1**).

It is very difficult to understand the regulation of endogenous condensates, which contain a large number of components and are regulated by complex pathways. In contrast, changes in phase separation of our bioorthogonal system are likely to be due to fundamental physicochemical changes in the cellular environment that will broadly impact phase separation in the cell. A predicted feature of a passive reporter of this cellular phase separation propensity is that the phase boundary can change without any change in the chemical nature of the system components or their interactions. Accordingly, we found that the phase boundary of our system can dramatically change when cells enter stationary phase without any change in apparent affinity between the two components as measured by FRAP (**Figure 2**).

During acute glucose starvation and stationary phase, the yeast cytoplasm undergoes dramatic changes, with many mesoscale protein assemblies appearing (*15*, *18*). We find that these conditions enhance phase separation of our system. During acute glucose starvation, the intracellular pH decreases and this likely drives some of these global changes (*13*, *14*, *41*). However, acute glucose starvation without acidification also enhanced the phase separation of our system, indicating that pH is one, but not the only factor impacting phase separation in this condition. Furthermore, the range of concentrations at which phase separation of our system takes place is larger in stationary phase than in log-phase, suggesting a wider heterogeneity of cells’ intracellular milieu in the former condition. Such heterogeneity might be associated with a bet-hedging strategy in stationary phase (*42*), whereby cells adjust the properties of their cytoplasm during stationary phase in diverse ways (**Figure 2**).

We introduced the system to the yeast KO collection (*27*, *28*, *43*) and measured its phase diagram in 2888 deletion backgrounds. We found 68 KOs that robustly alter phase separation of our synthetic system. These deletion strains exhibit an extraordinarily high number of phenotypes, as annotated in YeastMine (*33*). Phase separation has been implicated in a multitude of fundamental cellular processes, including transcription (*44*), signaling (*45–47*), enzyme activity (*48–50*), and stress adaptation (*51*). The recent explosion of biological processes associated with phase separation has led to considerable skepticism about the field (*52*). Our results suggest that phase separation may indeed make fundamental contributions to many aspects of biology.

One of the most frequent phenotypes among the outliers is *“increased heat sensitivity”*. Given that stress granules form in response to heat, and their formation through condensation is crucial for heat tolerance (*51*), it is compelling to speculate that the KOs with changed phase separation show increased heat sensitivity due to impaired stress granule assembly. A number of other phenotypes over-represented among the outliers of our screen relate to decreased resistance to DNA-damaging agents, including dieldrin, nitrosodiethylamine, and idarubicin (Supplementary Table 2). This is notable, because the DNA damage response involves phase separation (*53*), and altering phase separation globally could therefore render cells more vulnerable to DNA damage.

KOs with altered phase separation are enriched in TORC1 signaling-related genes. Furthermore TORC1 inhibition and nitrogen starvation alter phase separation of our system in wild type cells (**Figure 3-4**). This is in line with previous reports showing that changes in ribosome concentrations downstream of TORC1 signaling lead to changes in macromolecular crowding, which tunes protein phase separation (*9*). We used GEM diffusivity measurements to infer macromolecular crowding at the 40 nm length-scale to test whether the altered phase separation of a selected set of 6 KOs could be explained by this mechanism. ΔLST7 showed significantly higher GEM diffusivity, indicating lower ribosome density. This is in line with LST7 being necessary for TORC1 activation and thereby TORC1-mediated ribosome biogenesis. However, the other three strains did not show altered GEM diffusivity, implying that changes in macromolecular crowding is only one of multiple mechanisms by which TORC1 signaling can globally change phase separation of proteins.

We next measured diffusivity at the mesoscale in the same six KO strains, by tracking the motion of the condensates themselves, which are typically several hundred nanometers in size. We found that five out of six strains with changed phase separation also showed altered mesoscale diffusion. The process of phase separation occurs at multiple length-scales: nanoscale proteins condense into small droplets that then typically grow by diffusion, collision and coalescence (*54*). The cytoplasm is structured at various length-scales by ribosomes, polysomes, membranes and other unknown factors, each of which could frustrate condensate growth . This frustration of motion can be partly overcome by non-thermal active processes (e.g. actomyosin contractility (*55*, *56*)). We speculate that some mutants could affect the mesoscale structure or non-thermal motion of the cytoplasm at various length-scales (*56*, *57*).

Metabolomic profiling of these six strains showed alterations in nucleotide metabolism. This could reflect the overall energy state of the cell, influencing phase separation of our synthetic system indirectly. For example, active processes and ATP hydrolysis have been proposed to influence phase separation (*58*). However, nucleotide triphosphates could also act by decreasing the elasticity of the cytoplasm (*56*), or solubilizing the protein components (*59*), as ATP can act as a biological hydrotrope (*7*). In addition, tryptophan and related metabolites (Hydroxy-kynurenine, Kynurenate & Kynurenine) changed significantly. In line with this observation, ΔARO1, which is auxotroph for Trp, was among the high-confidence outliers of our screen. Moreover, proteomics data showed differential abundance of two genes related to Trp metabolism (ARO9 and BNA5) in those strains (Figure S7), and the Wikipathway “Tryptophan degradation” was enriched in the analysis of proteomics data reported in (*38*). The quality of the solvent in the cell has been shown to significantly affect biological assemblies (*20*), and water potential was recently shown to be intricately linked to phase separation (*60*, *61*). We speculate that certain metabolites including tryptophan and ATP might globally impact phase separation through change in solvent quality and water potential.

Taken together, our findings underscore that the propensity for intracellular protein phase separation is an emergent property influenced by a confluence of multiple pathways and factors. In classic genetic works, phenotypes such as adenine auxotrophy are directly and predictably influenced by specific genes (*62*). By contrast, we observed that metabolism (in particular amino acid, carbon, and nucleotide metabolism), energy state (e.g., NMP and NTP levels), pH, crowding, and additional factors such as water potential (*20*, *60*) or cell size (*63*), can collectively be associated with changes in phase separation. This complexity illustrates the intricacies of biological systems, where certain processes are controlled by well-defined pathways or modules (*64*), whereas others are governed by a complex landscape of interconnected factors (*38*, *65–68*). We provide a survey of this landscape as a step towards a systems-level understanding of phase separation homeostasis, to ultimately modulate it by targeted therapies.

## Acknowledgments

We thank Maya Schuldiner for several helpful discussions and for providing the strain YMS721. We thank Simon Alberti, Allan Drummond, members of E.D.L’s lab for helpful discussions. We thank Esther Weindling for technical support and Harry Greenblatt for IT support. We thank Alon Savidor, Corine Katina, and Hila Levy for carrying out proteomics profiling, and Sergey Malitsky and Maxim Itkin for carrying out the metabolomic profiling.

## Funding

E.D.L. acknowledges support from the European Research Council (ERC) under the European Union’s Horizon 2020 research and innovation program (grant agreement No. 819318), by the Human Frontiers Science Program Organization (Ref. RGP0016/2022), by the Israel Science Foundation (grant no. 1452/18), and by the Abisch-Frenkel Foundation. M.H. acknowledges support by the Pearlman Grant for student-initiated research in chemistry. L.J.H. was funded by NIH R01 GM132447, R37 CA240765, the NIH Director’s Transformative Research Award TR01 NS127186, and the Human Frontier Science Program (RGP0016/2022-102). Y.X. was funded by NIH R01GM107466.

## Methods

### Plasmids and strains

Throughout this work, BY4742/BY4743 (*69*) cells were used. For phase diagrams of treatments and conditions, the strain yMH07 was used. This strain is based on BY4743 diploids, and carries the plasmids encoding the synthetic system with 9.3×10^-6^ M affinity, selectable with hygromycin and G418. For the synthetic genetic array, the tetramer-encoding plasmid pMH01 was altered to replace the G418 resistance cassette with a resistance cassette to nourseothricin (Nat), using PIPE cloning (*70*). In short, primers binding in the *A. gossypii* TEF promoter and terminator that flank both the Nat and G418 resistance cassettes, were designed. Using these primers, the Nat resistance cassette, as well as the plasmid backbone (pMH01, (*23*)) were amplified. Then, the products of the two PCRs were used to extend and amplify each other in a third PCR reaction, and the product was transformed into competent DH5α *E. coli* cells via heat shock. The plasmid was purified (Promega PureYield™), and verified by sequencing of the new resistance cassette. To generate the library, this new plasmid was co-transformed with pMH06 (the dimer with 9.3×10^-6^ M affinity) into the SGA donor strain YMS721 (*71*). The yKO library was obtained from the bacteriology unit of the Weizmann Institute of Science. For the validation of the nitrogen starvation, BY4743 cells expressing either a second synthetic system (*22*) or EDC3-mTagBFP2 were used. We mated BY4742 cells expressing the ferritin dodecamerization domain fused to sfGFP and the intrinsically disordered region of hnRNP with wild type BY4741 cells and selected for diploids using SD -M-K agar plates. The plasmid encoding EDC3-mTagBFP2 was synthesized (Twist Bioscience) and transformed to BY4743 cells using heat shock. The plasmids and strains used and generated in this study are summarized in Supplementary Table 3.

### Library creation via synthetic genetic array

An SGA with the KO/Phase SGA donor strain and the yeast deletion collection was performed according to (*29*). In short, the donor strain (Hygromycin and Nourseothricin selection) and the yeast KO collection (*28*, *43*) (G418 selection) were printed on YPD agar plates with the respective antibiotic, using a robotic liquid handling system (Tecan Evo 200). The donor strain was mated to the library, by transferring cells onto YPD agarose plates containing no selection, followed by a 24h incubation at 30 °C. Subsequently, diploid selection was conducted twice, by transferring the colonies to YPD plates containing G418, nourseothricin, and Hygromycin. The diploids were then driven to sporulation and subsequently haploids were selected (SD-LRK+thialysine+canavanine), followed by a final haploid selection (final selection: SD-LRK+thialysine+canavanine+G418+nourseothricin+Hygromycin). The final library was transferred to a liquid culture in 384-well plates with 35 µL final selection media. After three days of growth, 15 µL glycerol were added to a final concentration of 15% and cells were stored at -80 °C. Prior to imaging, cells were transferred to polypropylene 384-well plates (Greiner™) containing 35 µL selection media, and grown for three days. From this culture, 1 µL was transferred to 35 µL fresh final selection media in optical glass bottom 384-well plates (Greiner™) and cells were grown for 6 h at 30 °C prior to imaging.

### Cell culture and treatments

For log-phase cultures, yMH06 (*23*) cells, *i.e.* BY4743 (*69*) with the synthetic system, were cultured for three days, diluted to an OD_600_ of 0.05-0.1, and grown for at least 6 h. For the hyperosmotic shock, cells were grown for 6 h in an ConA-coated optical well, and imaged. Then, sorbitol (Sigma) was added to a final concentration of 1M and cells were either followed by time-lapse microscopy, or phase diagrams were measured 15 min and 1h after the addition. For the stationary phase phase diagram and FRAP experiment, cells were grown for a week to generate large enough condensates. Cells were then grown to logarithmic phase for 6 h, and subsequently incubated at 30 °C for three days without dilution and agitation prior imaging. For acute glucose starvation, cells were washed twice into SD media lacking glucose, where the lack of glucose was osmotically balanced with sorbitol. The time between the first wash into glucose free media and imaging was 15 min. Re-addition of glucose was conducted 15 min after the first wash into glucose-free media. For glucose starvation at pH7.5, cells were washed into glucose-free or regular SD media that was adjusted to pH7.5 using KOH.

For Rapamycin treatment, 1 µL of logarithmically growing cells was transferred into 35 µL SD media in an optical glass bottom 384-well plate (Greiner™) and 4 µL Rapamycin (BioShop) were added to yield a final concentration of 100 nM. As a control, DMSO (Sigma-Aldrich) was diluted with SD media in the same manner as the Rapamycin stock solution, and 4 µL were added to the control wells. For nitrogen starvation, log-phase cells were washed twice with PBS, resuspended in synthetic defined media lacking amino acids and ammonium sulfate, and incubated for 6 h at 30 °C. For amino acid and nitrogen starvation, yMH06 cells, cells that expressed the second synthetic system, or the EDC3-BFP construct were washed twice into media lacking amino acids and ammonium sulfate. Imaging was conducted 6h later. As a control, cells were washed into complete media at the same time point, 6h prior imaging.

### Imaging, image processing and FRAP

Imaging and FRAP experiments were conducted as described in (*23*). For automatic cell segmentation, we used a custom deep learning algorithm. The network was based on the UNET architecture (*72*). The input for training was the 2048×2048 pixels brightfield images, and the ground truth was based on images of cells expressing a cytosolic mNeonGreen fluorescent protein. The segmentation of the fluorescent images was achieved with YeastSpotter (*73*), and the binary masks generated formed the basis of the ground truth. The ground truth images were stacks consisting of 9 slices separated by 1 micron z-steps. We only considered the masks of the center slice where the fluorescence intensity was maximal (i.e., where the total fluorescence within the mask decreased from slices 4 to 1 and from 6 to 9). This procedure aimed at forcing the network to identify cells in focus. A Pytorch implementation of the network was trained on these images and used to identify cells in brightfield images. The ROIs so identified were subsequently processed as in (*23*, *74*).

### Measuring phase diagrams

To measure phase diagrams, data was processed using custom scripts in R. To determine whether a cell contains a condensate, each cell pixels’ intensities were divided into 10 bins. A cell was deemed to have a condensate, if it had a minimum of 600 A.U. GFP and 300 A.U. RFP intensity in their brightest intensity bin, and if the brightest pixel was at least 5 times (GFP) or 2.5 times (RFP) brighter than the median intensity value. Median RFP and GFP intensities were converted to concentrations using the same approach as in (*23*). Finally, RFP and GFP concentrations were plotted against each other. In order to identify and fit the phase boundaries, we selected cells containing condensates as the concentrations of free dimer and tetramer are reflecting the phase boundaries. We note that this approach differs from the previous one (*23*), where we considered the excluded region to visualize (but not quantitatively analyze) the phase boundary. Indeed, the previous approach (i.e., identify a region with an absence of data points) was not amenable to a fitting procedure. Furthermore, we note that not all cells containing condensates are detected, as certain condensates fall below the detection limit due to their small size. However, the cells included in our analysis unequivocally exhibit condensates, enabling the delineation of the phase boundary.

### Measuring phase diagrams in high-throughput

To measure the phase diagram of the library, two methods were employed, which we refer to as the ‘center method’ and ‘branch method’, respectively. For both methods, only strains with more than 10 cells with condensates were considered.

#### Center Method

For the first method to analyze the library’s phase diagrams, the ‘fel’ function with the ‘direct’ method (least squares fit) of the package “Hysteresis” for R was used on the decimal logarithm of the concentration values of tetramer and dimer (*75*). From the resulting estimates, the center points were recorded for each strain’s phase diagram’s fit (**Figure 3**). To select outliers for further validation, z-scores were calculated for both coordinates, using the ‘scores’ function of the ‘outlier’ package in R (*76*) and strains that had a lager absolute z-value of 2 (RFP) or 2.3 (GFP) were selected for further validation.

#### Branch Method

For the second method of analysis, a reference phase diagram of random cells containing condensates in a plate was generated. Next, the decimal logarithm of the cytoplasmic concentrations was calculated, an ellipse was fitted using the ‘fitConic’ package in R (*77*) and the line passing through this ellipse’s center with a slope of 1 was used to split the data into the ‘upper’ and ‘lower’ branches (Supplementary Figure 1). Data points closer to the diagonal than 0.1 were excluded. Then, a polynomial regression was used to fit each of the two branches, representing average boundaries for strains in a full multiwell plate. We then analyze each strain relative to these average boundaries. Specifically, we measured the fraction of cells containing puncta inside the upper and lower branch of the background. Outliers were selected as in the center method.

To validate the outliers, 301 strains that were outliers in the screen were transferred to a new 384-well plate, alongside with 11 reference strains that did not show a changed phase diagram, where one strain (ΔCTA1) was placed in 29 positions across the plate to control for position effects. The results of the validation screens were normalized for expression, where expression was estimated as the median concentration of cells with either 100 times more GFP than RFP concentration, or vice versa (Supplementary Figure 6). Strains were deemed high confidence outliers when the normalized measurements were significantly different from reference strains in at least two out of three replicates (α=0.05, two-sided Z-test with Bonferroni-Hochberg *P*-value adjustment, Supplementary Figure 6).

### GO-term and phenotype analysis

GO-term analysis (*31*, *32*) was conducted using gProfiler (*36*) with Benjamini-Hochberg (BH) *P*-value adjustment for multiple testing, and the 2,888 KOs screened in our library formed the background against which enrichment was calculated. Only GO-terms highlighted by gProfiler were considered. For analyzing the outlier’s phenotypes, a table with all phenotypes for all genes was retrieved from YeastMine (*33*), https://yeastmine.yeastgenome.org/yeastmine/service/template/results?name=Phenotype_Tab_New&format=tab&size=10, 28/3/22). This data was subsetted for phenotypes of null mutants of S288C strain that were sampled in the library, and the number of unique phenotypes was counted for each of the 68 outlier and 11 reference strains. For comparison, 100 sets of 68 random KOs were sampled from all screened strains, and the median number of unique phenotypes was calculated for each set. For phenotype enrichment, the average and standard deviation of the frequency of each phenotype in 100 sets of 68 random KO strains was calculated. This random frequency was compared to the frequency of each phenotype among the 68 outliers, using Z-tests with BH *P*-value adjustment. A phenotype was considered enriched if *P*(adjusted)<0.05 in at least two high confidence outliers. In **Figure 3H**, only the top ten phenotypes in terms of their frequency among the high confidence outliers are shown, ordered by their adjusted *P*-value. Supplementary Table 2 contains all significantly enriched phenotypes.

### GEM diffusivity

*Saccharomyces cerevisiae* knockout strains were revived from −80°C freezer on YPD plates for overnight growth. To integrate 40nm-GEMs into the yeast strains, plasmid pLH497 (pINO4-pfv-Sapphire::Leu2) was linearized by restriction enzyme SnaBI, and yeast transformation was performed using heat shock. Positive colonies were selected by the SC-Leu2 plate and confirmed by visible 40nm-GEM particles under HILO TIRF microscope.

To perform 40nm-GEM’s tracking experiment, a patch of yeast cells were cultured for three days in 5-mL synthetic complete media with 2% glucose, 0.1% monosodium glutamate and 100 μg/ml G418. And cells were refreshed at O.D.∼0.1 for 6 hours before applying them on 1mg/ml concanavalin A (Alfa Aesar, Cat. no. J61221) coated 384-well (Cellvis, Cat. no. P384-1.5H-Nor) glass bottom imaging plates. GEM movies were captured by TIRF Nikon TI Eclipse microscope in highly inclined thin illumination mode (HILO), using 488 laser for excitation with 100% power. Fluorescence was recorded with a sCMOS camera (Zyla, Andor) with a 100x Phase, Nikon, oil NA = 1.4 objective lens (pixel size: 0.093 μm). Cells were imaged at 100 Hz (10 ms per frame) for a total of 4 sec.

The tracking of particles was performed with the Mosaic suite of FIJI using the following parameters: radius = 3, cutoff = 0, Per/Abs: variable, a link range of 1, and a maximum displacement of 7 px, assuming Brownian dynamics. Individual yeast cells were segmented by YeastSpotter (*73*). And trajectories within individual cells were then analyzed by GemSpa that we developed in house: https://github.com/liamholtlab/GEMspa/releases/tag/v0.11-beta. Mean-square-displacement (MSD) was calculated for every 2D trajectory, and trajectories continuously followed for more than 11 time points were used to fit with linear time dependence based on the first 10 time intervals to quantify time-averaged MSD: MSD(T) = 4D_eff_T, where T is the imaging time interval and D_eff_ is the effective diffusion coefficient with the unit of μm^2^/s.

### Condensate diffusion

Cells were grown in library media for three days, before being inoculated at ∼OD 0.1 and grown without agitation for 6h to log phase. After centrifugation and removal of supernatant, 1uL concentrated cell solution was placed onto a 384 well plate and spread gently using a pipette tip. 50ul of 1% low melting temperature agarose (Sigma), was pipetted onto cells. Time lapses of droplets movement within cells were recorded with images taken every 2s for 10min. For each time point, 5 Z-stacks with a step size of 1.8μm were taken. Finally, a maximum intensity projection of the Z-stacks was generated using Fiji (*78*). Additionally, binary mask files were generated from brightfield images of yeast cells to enable the identification of individual cells. Droplets were identified using the Laplacian of Gaussian filter in the Trackmate plugin in Fiji, with a radius of 0.75 μm and varying thresholds (*79*). Subsequently, droplet trajectories were determined through the Simple Linear Assignment Problem (LAP) tracker, which linked the identified droplets using the following parameters: Linking max distance = 0.6 μm, Gap-closing max distance = 0.6 μm and Gap-closing max frame gap = 1. Only droplet trajectories with at least 10 time points were used to calculate their effective diffusivities. This involved fitting the first 10 time intervals of their mean squared displacement using custom MATLAB code. Droplets were uniquely identified within each cellular region of interest (ROI) based on their locations. The droplet diffusivities within each individual cell were averaged, and the resulting values were plotted for each mutant condition and compared with those of the ΔHIS3 control.

### Metabolomics

#### Sample Preparation and LC-MS

Extraction and analysis of polar metabolites was performed as previously described in (*80*, *81*) with the following modifications: yeast cell pellets were extracted with 1 ml of a pre-cooled (−20°C) homogenous methanol:methyl-tert-butyl-ether (MTBE) 1:3 (v/v) mixture, The tubes were vortexed and then sonicated for 30 min in an ice-cold sonication bath (taken for a brief vortex every 10 min). Then, double deionized water : methanol (3:1, v/v) solution (0.5 mL) containing internal standard: C13 and N15 labeled amino acids standard mix (Sigma, 767964) was added to the tubes followed by centrifugation. The organic phase was transferred into a 2 mL Eppendorf tube. The polar phase was re-extracted as described above, with 0.5 mL of MTBE, and the organic phase was combined with the first extraction. The polar phase was moved to a new Eppendorf, dried for 1.0 hr by N_2_ to remove organic solvents, and then lyophilized. Before the injection into the LC-MS, the polar phase sample pellets were dissolved using 150 uL DDW-methanol (1:1), centrifuged twice (13,000 rpm) to remove possible precipitants, and transferred to an HPLC vial.

Metabolic profiling of the polar phase was done as described in (*81*) with modifications described below. Briefly, analysis was performed using Acquity I class UPLC System combined with mass spectrometer Q Exactive Plus Orbitrap™ (Thermo Fisher Scientific), which was operated in a negative ionization mode. The LC separation was done using the SeQuant Zic-pHilic (150 mm × 2.1 mm) with the SeQuant guard column (20 mm × 2.1 mm) (Merck). The Mobile phase B: acetonitrile and Mobile phase A: 20 mM ammonium carbonate with 0.1% ammonia hydroxide in DDW: acetonitrile (80:20, v/v). The flow rate was kept at 200 μl min−1, and the gradient was as follows: 0-2min 75% of B, 14 min 25% of B, 18 min 25% of B, 19 min 75% of B, for 4 min.

#### Data Pre-Processing and analysis

Data processing was done using Progenesis QI (Waters). The detected compounds were identified by accurate mass, retention time, isotope pattern, fragments, and verified using an in-house-generated mass spectra library. The results were normalized by relative abundances of internal standards and by OD.

Data analysis was done in R 4.1.1. The data were log-transformed and normalized by loess normalization using the NormalyzerDE package (*82*). Imputation was performed using the imputePCA function from missMDA(v1.18) package (*83*). All samples were considered as biological replicates. ΔHIS3 was used as a reference strain. BH was used for multiple testing corrections. A metabolite was considered changed if | log2FC | > 1 and *P*(adjusted)<0.05. Heatmaps were generated using the pheatmap package (*84*).

### Analysis of proteomics dataset

The abundance of each protein in the 62 high confidence outlier KO strains reported in (*38*) was divided by its mean abundance in the reference strains and ΔHIS3. There were 41 proteins exhibiting ≥2-fold change in at least six high confidence outlier strains, and the abundance of those proteins across the 62 KO strains gave a matrix subsequently used for clustering (**Figure 5F**) using the ‘hclust’ function in R with Ward distance. GO-term enrichment analysis was then performed on each cluster, using gProfiler (*36*). When conducting the GO-term enrichment, we used all detected proteins (n=1850) as the universe, and BH *P*-value adjustment to correct for multiple testing.

## Supplementary Figures

**Supplementary Figure 1:**
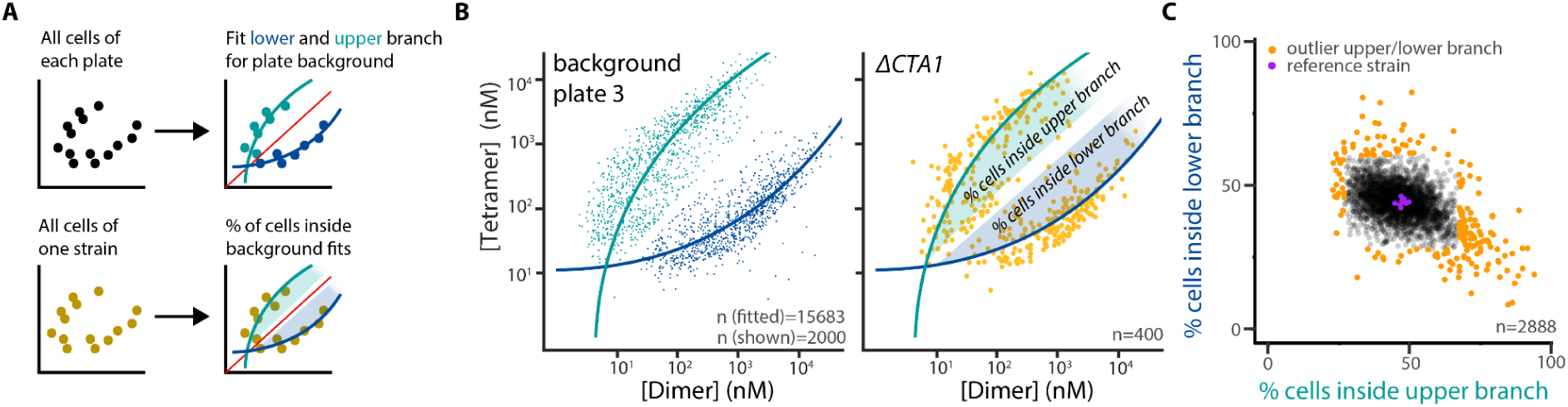
The branch method for detecting outliers. **A.** Schematic of the branch-method of analysis. First, a reference fit is generated from all strains present in the same 384-well plate, by dividing the data of cells with condensates by the diagonal, and fitting two polynomial regressions for the “upper” and “lower” branches of the reference phase boundary. Then, each strain’s phase diagram is compared to the background fit of its plate. The fraction of cells inside the “upper” and “lower” branches is recorded. **B.** Background fit for plate 3. We considered separately cells falling on the upper- (cyan) or lower- (darker blue) branch of the phase boundary. Polynomial regressions were fitted to both branches. Phase diagram of ΔCTA1 strain compared to the reference fit of plate 3 is shown on the right. **C.** Result of the branch method analysis for all strains in the screen. The percentage of cells inside the lower branch is plotted against the percentage of cells inside the upper branch. Purple points represent reference strains, orange points are outliers selected for further validation from this method.

**Supplementary Figure 2:**
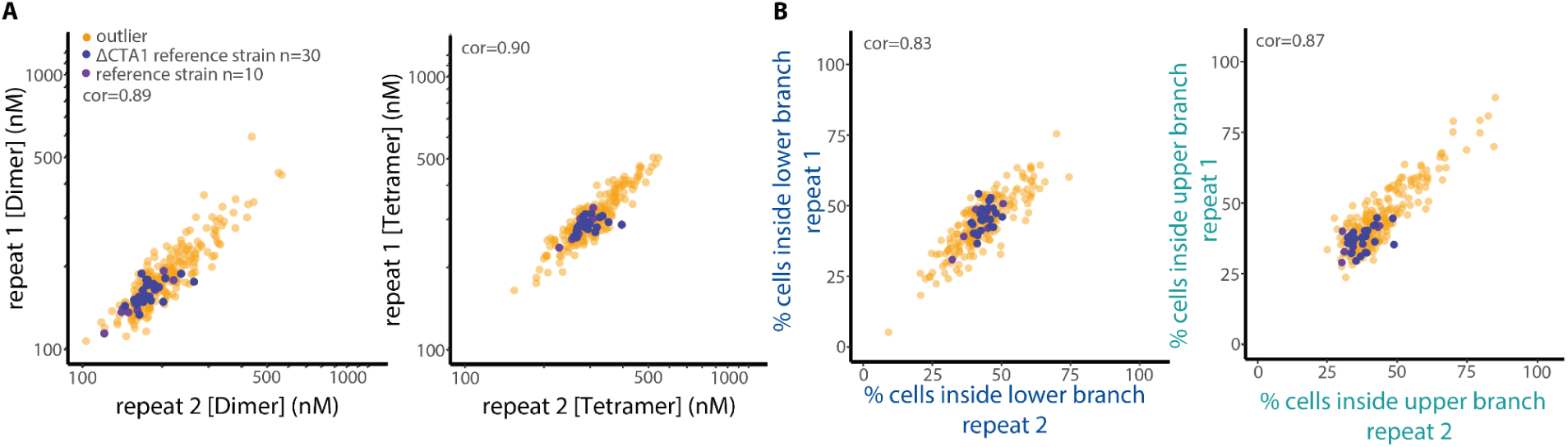
The results of the validation screens correlate well. **A.** Results of the first validation screen are plotted against the results of the second one for a. The center method’s x- (left) and y-coordinate (right), and **B.** the branch method. Pearson correlation coefficients are indicated above. Outlier strains from the full screen are shown in orange and purple denote reference strains, where ΔCTA1, repeated 30 times in the plate, is represented in dark purple.

**Supplementary Figure 3:**
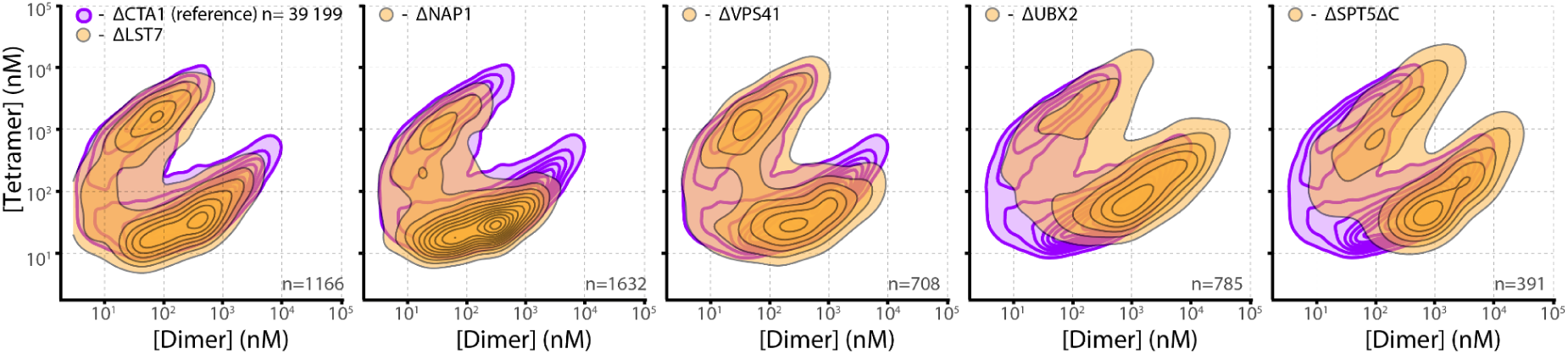
Phase diagrams of five outliers of our screen. Purple lines show the 2D density of the reference strain ΔCTA1. The same reference strain data is shown across all diagrams. Gray lines and orange areas show the outliers as indicated above. The number of cells is indicated in the inset.

**Supplementary Figure 4:**
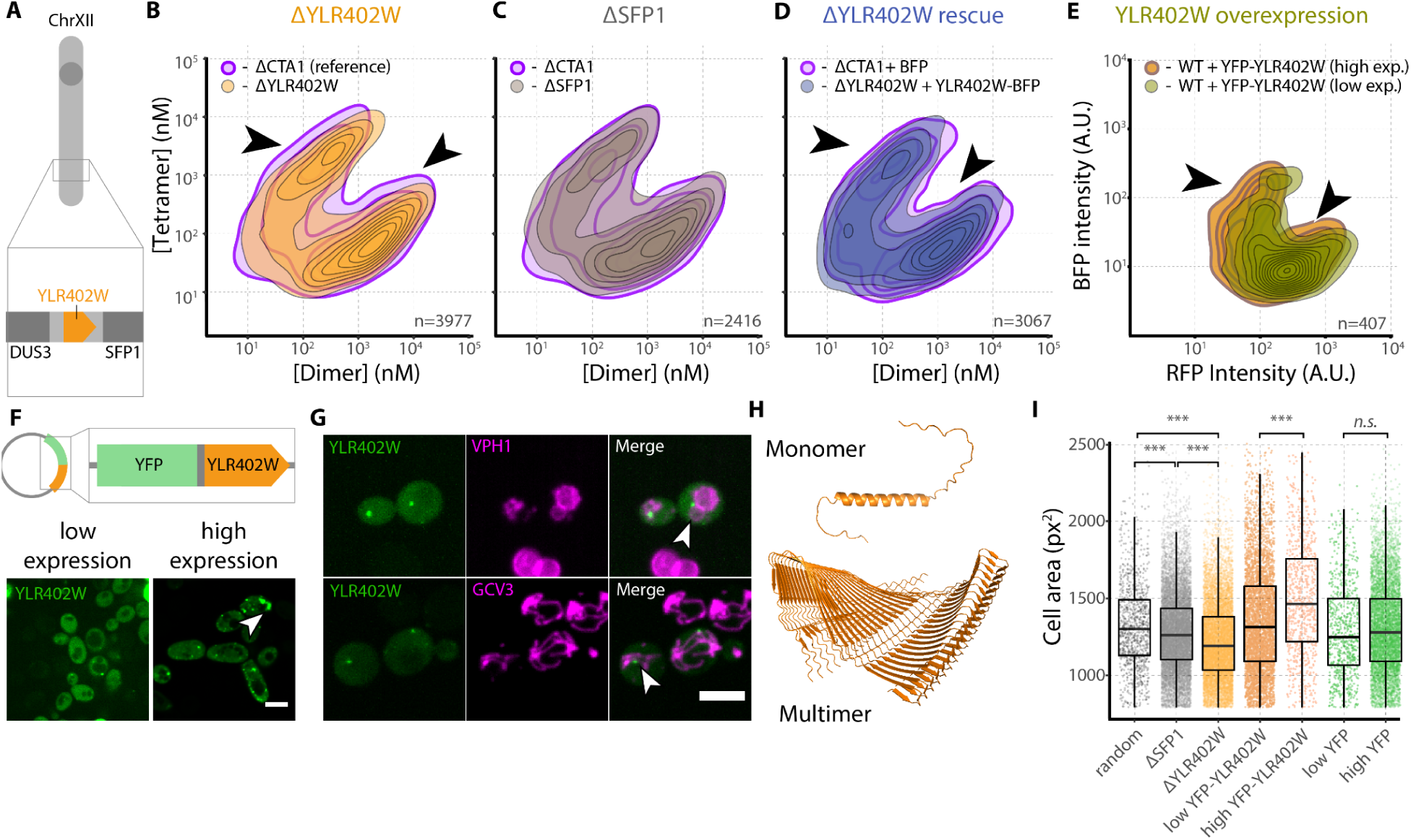
YLR402W is an open reading frame of unknown function that influences phase separation of the synthetic system, as well as cell size. **A. Schematic view of YLR402W genomic context** on Chromosome XII, between DUS3 and SFP1 ORFs. **B. ΔYLR402W changes the phase diagram of our system.** Orange area represents the 2D density of the dilute phase concentrations of tetramer and dimer in cells with condensates of the ΔYLR402W strain (n=3977). The purple area represents the 2D density of ΔCTA1 cells, as a reference. Black arrows highlight positions where the phase diagram of ΔYLR204W deviates from the reference. **C. ΔSFP1 does not change the phase diagram of our system.** Gray area represents the 2D density of the data of ΔSFP1 cells. The purple area represents the 2D density of the ΔCTA1 reference strain (same data as in b is shown). **D. The phase separation phenotype of ΔYLR402W can be rescued by expressing YLR402W from a plasmid.** The blue area shows the 2D density of ΔYLR402W cells expressing BFP-tagged YLR402W. The purple area represents the 2D density of ΔCTA1 cells expressing BFP alone, for comparison. **E. The phase diagram of wild type cells changes in dependence of YLR402W expression level.** Cells carrying a version of the synthetic system with a BFP-tagged tetramer were transformed with a plasmid encoding YFP-tagged YLR402W. The phase diagram was generated for cells with low YFP signal (green area), and high YFP signal (orange area). Low YLR402W expression changes the phase diagram similarly to ΔYLR402W. Arrows highlight areas where the phase boundary is shifted in a similar direction as in B. **F.** A schematic of the plasmid encoding Venus-YLR402W shown above. Representative micrographs of wild type cells expressing the construct at low (left) and high (right) concentrations are shown below. The white arrow highlights an elongated structure in a cell expressing high levels of YFP-YLR402W. Scale bar= 5 µm. **G.** The puncta observed in cells expressing low levels of YLR402W (green) localize adjacently to the vacuole and mitochondria (magenta) marked with VPH1-RFP (vacuole, upper panel) and GCN3-RFP (mitochondria, lower panel). The white arrows highlight the position of YLR402W puncta adjacent to the tagged organelles. Scale bar= 5 µm. **H.** The structure prediction of a monomer by AlphaFold2 (*85*, *86*) shows an alpha helix flanked by disordered regions. Interestingly, predicting a 20-mer instead of a monomer with AlphaFold2 suggests this protein can oligomerize through formation of a beta sheet. **I.** YLR402W may be involved in cell size regulation. The area in pixel square of automatically segmented cells in our micrographs was recorded and compared across strains. ΔYLR402W cells (bright orange, n=10166) are significantly smaller than randomly sampled cells from the same plate of the KO collection expressing our system (gray, n=1000, Welch’s two sample t-test *P*<10^-16^). ΔSFP1 cells (dark gray, n=10800) are also significantly smaller than the random KO cells (Welch’s two sample t-test *P*=2.4*10^-6^), yet ΔYLR402W cells are even smaller (Welch’s two sample t-test *P*<10^-16^). Cells with more than 200 A.U. of YFP intensity were deemed to express high YFP-YLR402W levels. These cells show an increased cell size (n=780), compared to cells expressing the construct at lower levels, *i.e.* with a maximum YFP intensity less than 200 A.U. (n=4331, Welch’s two sample t-test *P*<10^-16^). Cells that expressed YFP alone from a plasmid did not show a significantly altered cell size when expressing high (n=6815) or low levels (n=742, cells were divided into those with more/less than 200 A.U. YFP intensity in the brightest pixels, Welch’s two sample t-test *P* = .3). n.s.: not significant, ***:*P*<0.0001

**Supplementary Figure 5:**
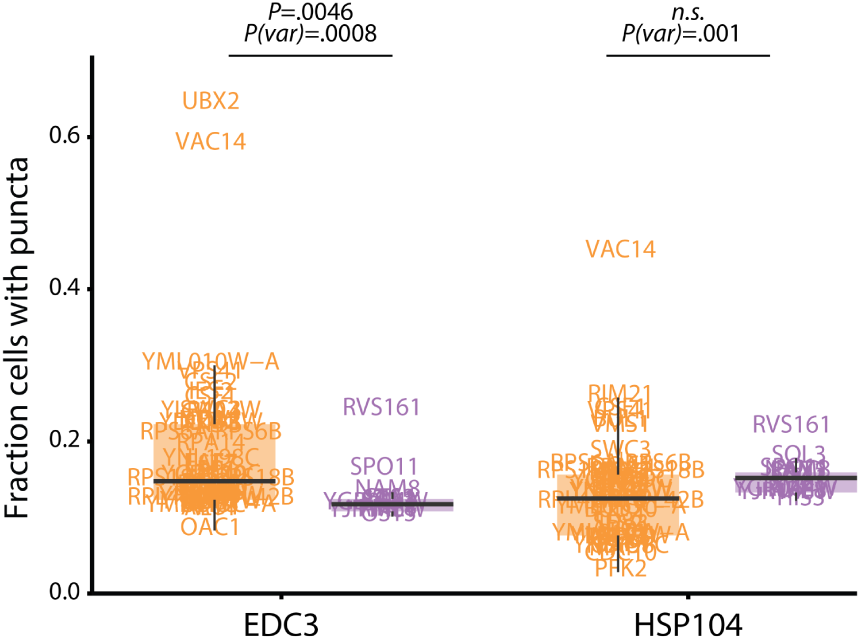
Impact of phase-separation influencing KOs on EDC3 and HSP104 localization. EDC3 (left) and HSP104 (right) were endogenously tagged with Scarlet-I in the genetic background of high-confidence KOs (orange), or in the genetic background of reference strains (purple). We imaged the strains and calculated the fraction of cells containing puncta for each (y-axis). Boxes delineate the first and third quartiles, the middle line corresponds to the median, the upper and lower whiskers extend to the highest and lowest values, and at most 1.5 times the interquartile range. *P*-values for Welch’s two-sample t-tests and F-tests are indicated above. Both ORFs were tagged in YMS721, using homologous recombination, and inserted to the genome of 46 KO strains identified to change phase separation, via SGA. We note that 22/68 high confidence outliers were not included in the SGA, because the definition of high confidence outliers changed after this experiment was performed, in order to control for changes in expression level (Figure S6).

**Supplementary Figure 6:**
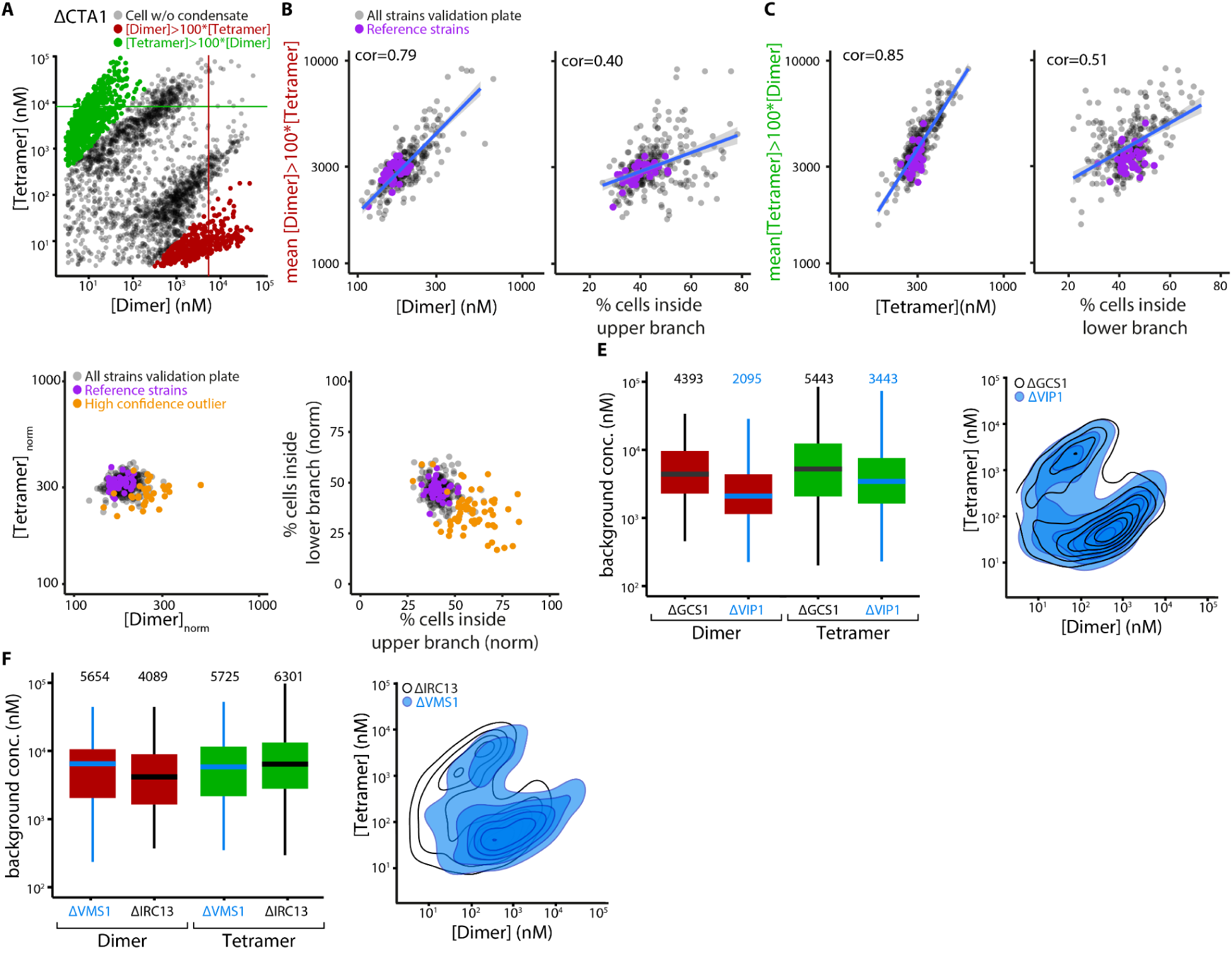
Normalization and picking of high confidence outliers from validation screen. **A.** Phase diagram of the reference strain ΔCTA1, with highlighted cells used for expression estimation. The dilute phase concentrations of tetramer and dimer of cells without condensates is plotted. Cells with 100 times more dimer than tetramer (red) and vice versa (green) were used to estimate the expression of the system in the KO strain. **B-C. Correlations of background expression of the system with the measurements of the center- and branch method of one validation screen.** B. The estimated dimer expression is plotted against the center point of the ellipse in the dimer dimension (left) and the percentage of cells inside the upper branch (right). C. The estimated tetramer expression is plotted against the center point of the ellipse in the tetramer dimension (left) and the percentage of cells inside the lower branch (right). Gray points represent all strains of the validation plate, purple points represent reference strains. The correlation is indicated in the inset. **D. Result of the validation screen with normalized measurements of the center method (left) and the branch method (right). E. Example of strains with different expression levels of the tetramer and dimer, but same phase diagrams. F. Example of two strains with similar expression, but different phase diagrams.**

**Supplementary Figure 7:**
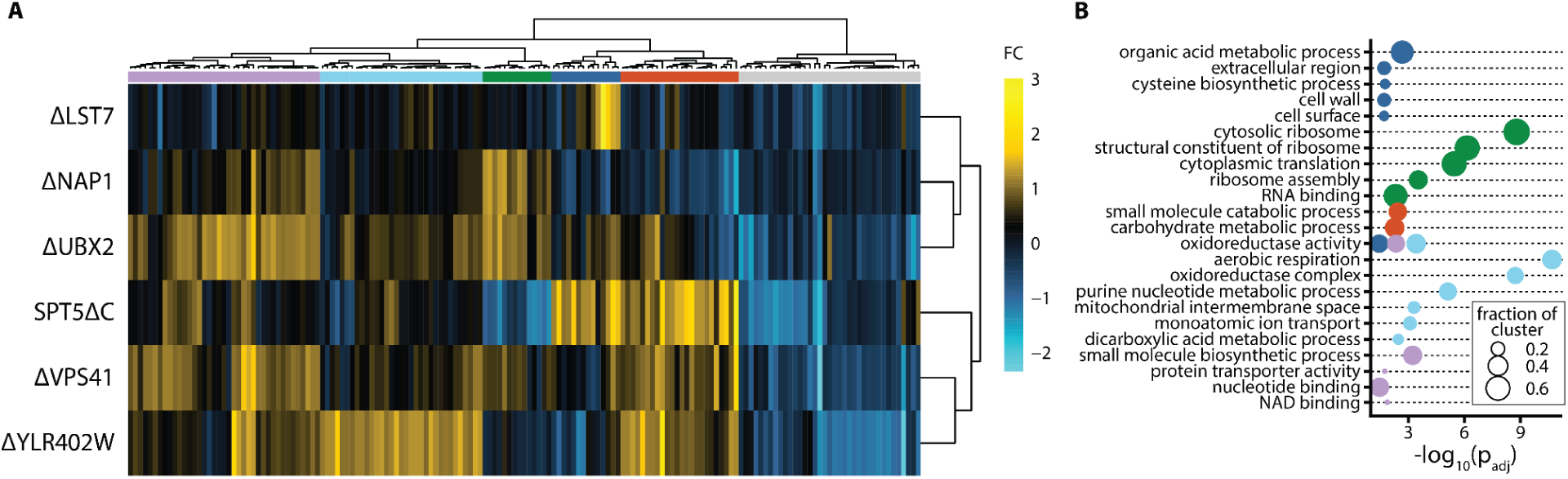
Proteomic profiling of the six selected strains. **A. Heatmap showing the fold-changes of proteins that are significantly changed in at least one strain, compared to ΔHIS3.** Six clusters were generated and are indicated as colors above. Cell pellets were subjected to lysis and in-solution tryptic digestion using the S-Trap method (by Protifi) (*87*, *88*) followed by a solid phase extraction (Oasis HLB) clean-up step. The resulting peptides were analyzed using nanoflow liquid chromatography (nanoAcquity) coupled to a mass spectrometer (Orbitrap Exploris 480). Each sample was analyzed on the instrument separately in a random order in discovery mode. Raw data were processed with MaxQuant v1.6.6.0. (*89*) The data were searched with the Andromeda search engine against a database containing protein sequences of Saccharomyces cerevisiae as downloaded from UNIPROT (*90*), and appended with common lab protein contaminants. The following modifications were defined for the search: Fixed modification-cysteine carbamidomethylation. Variable modifications-methionine oxidation and N-terminal protein acetylation. Decoy hits were filtered out as well as common lab contaminants. Peptide- and protein-level false discovery rates (FDRs) were filtered to 1% using the target-decoy strategy (*91*). Data analysis was done in R 4.1.1. Only proteins that were identified in 5 out of 6 replicates were kept for further analyses. Proteins that had only one unique peptide across all samples were discarded. All lab contaminants, recombinant proteins (antibiotics) and reverse sequences were removed before analysis. The data were log-transformed and normalized by loess normalization using the NormalyzerDE package (*82*). To minimize information loss on account of NA values (peptide intensities below detection limit), imputation was performed using the imp4p package (*92*). In this package, the “pca” algorithm was used for imputation of values missing completely at random (MCAR) and “igcda” for values missing not at random (MNAR). The Limma package (*93*) was used for identifying differentially regulated proteins. Here, all repeats were considered independent, therefore there were six samples per strain. Benjamini Hochberg correction was used to correct for multiple testing. A protein was considered differentially expressed if | log2FC | > 1 and *P*(adjusted) <.05, as reported by Limma. Heatmap of differentially expressed proteins was generated using the pheatmap package (*84*). Clustering into six clusters was done using the ‘hclust’ function in R. **B. GO-term analysis of the proteins that are differentially expressed in each of the six clusters.** The analysis was conducted using gProfiler (*36*), *P*-value adjustment was done using Benjamini-Hochberg correction. Only GO-terms associated with at least 10% the proteins in the cluster and that were highlighted by gProfiler default algorithm are shown. The size of the dot represents the fractions of proteins in each cluster associated with the respective GO-term. The clusters are enriched in several GO-terms, including “*aerobic respiration”, “organic acid metabolic process”, “small molecule catabolic process”, “small molecule biosynthetic process”, “cytosolic ribosome”*, or *”oxidoreductase complex”*.

**Supplementary Table 1:**
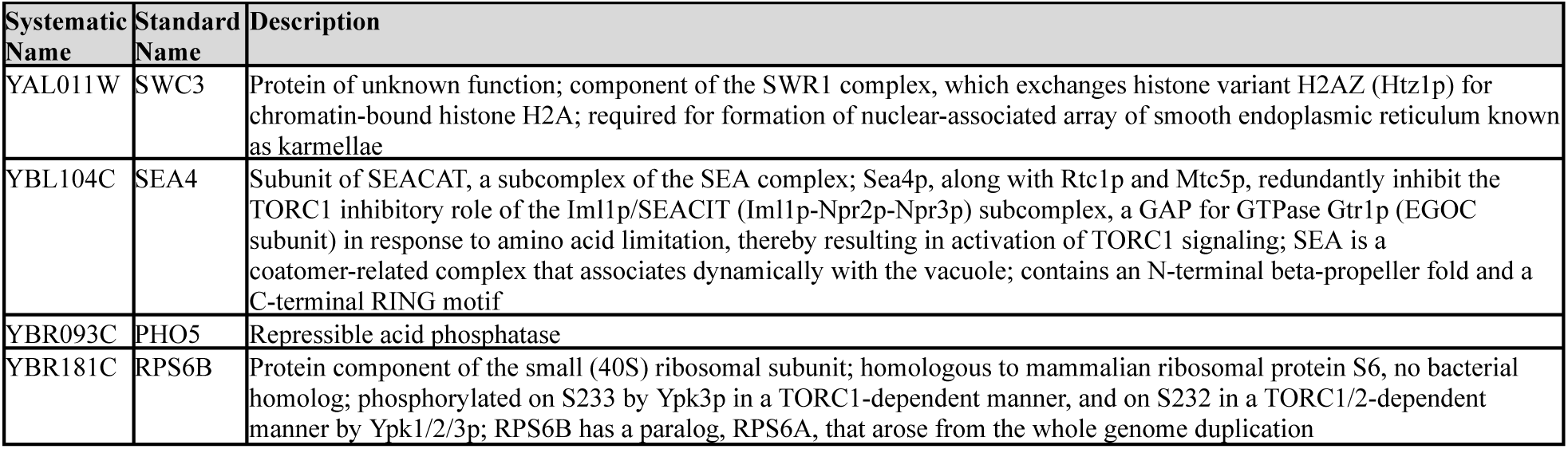
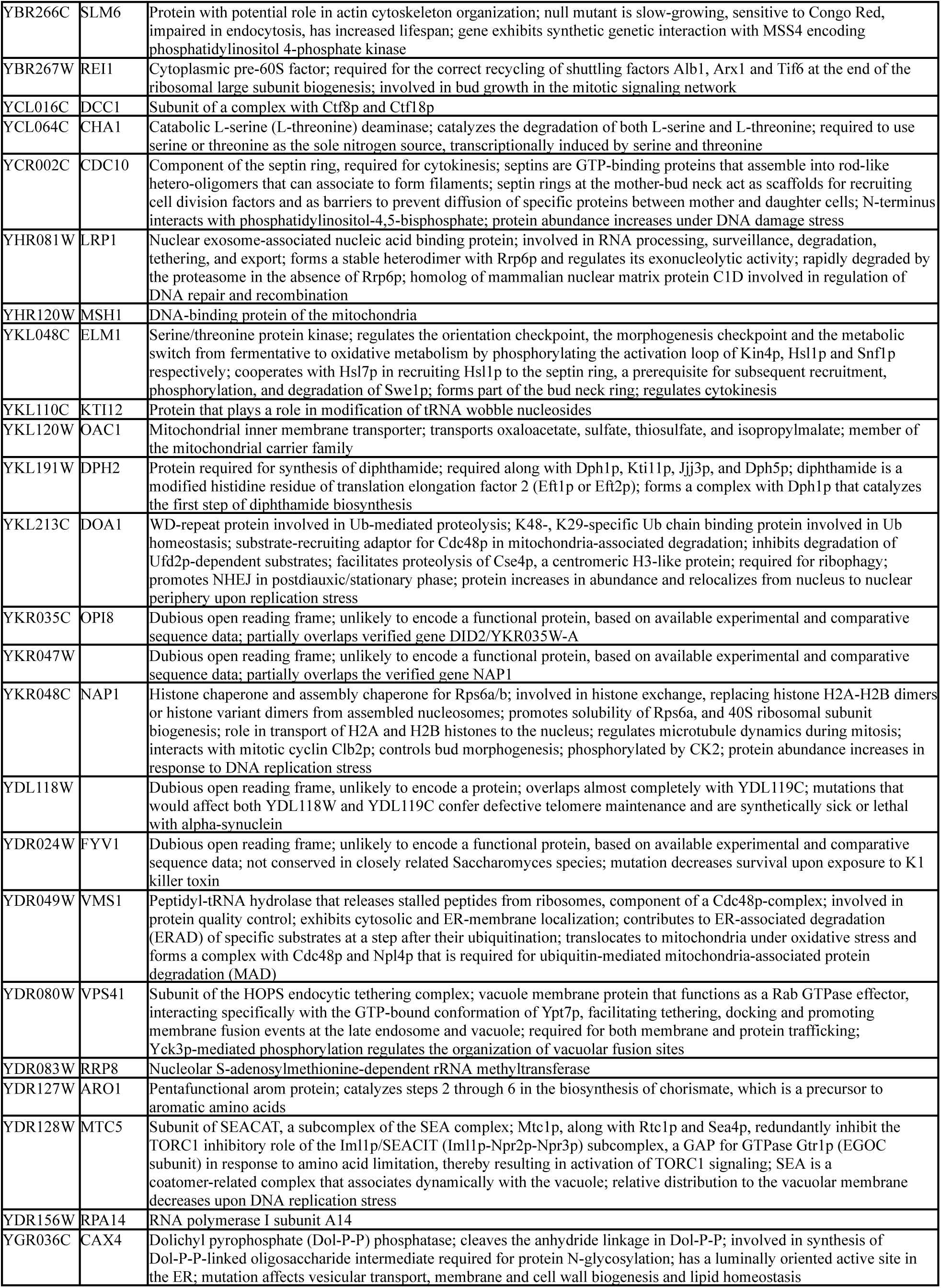
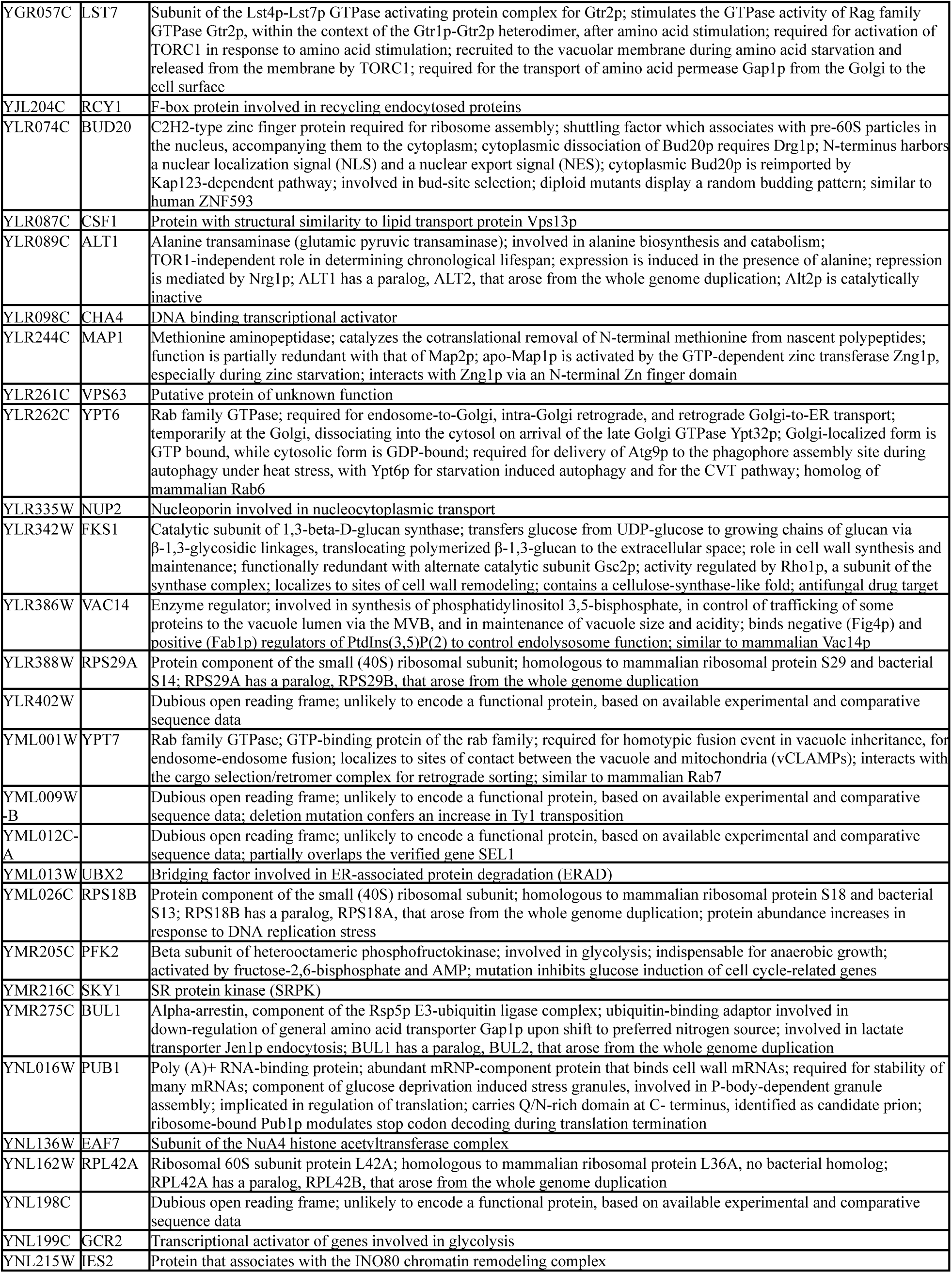
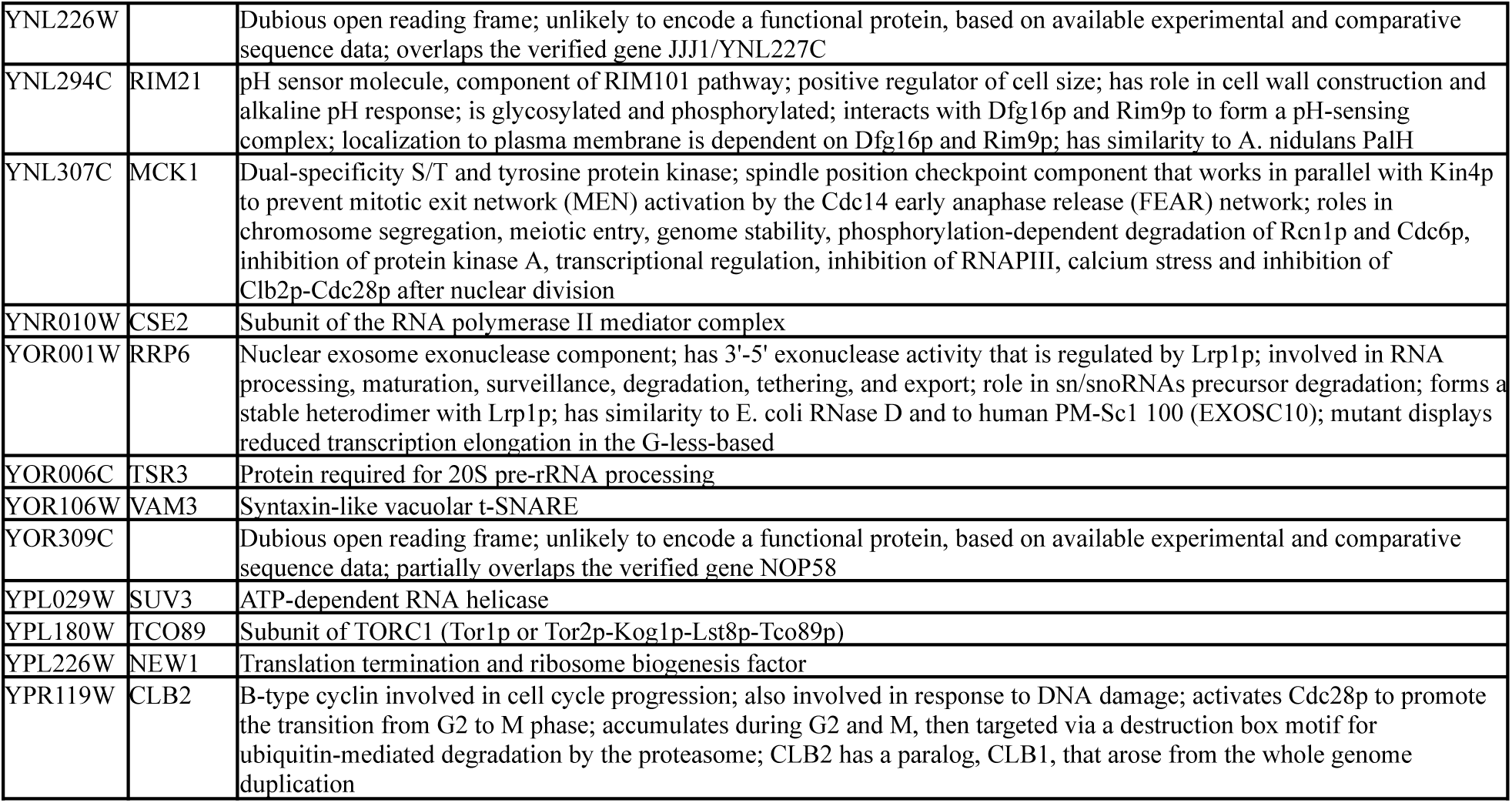
All high confidence outliers of the screen and their annotations as in the SGD (*94*).

| Systematic Name | Standard Name | Description |
| --- | --- | --- |
| YAL011W | SWC3 | Protein of unknown function; component of the SWR1 complex, which exchanges histone variant H2AZ (Htz1p) for chromatin-bound histone H2A; required for formation of nuclear-associated array of smooth endoplasmic reticulum known as *karmellae* |
| YBL104C | SEA4 | Subunit of SEACAT, a subcomplex of the SEA complex; Sea4p, along with Rtc1p and Mtc5p, redundantly inhibit the TORC1 inhibitory role of the Iml1p/SEACIT (Iml1p-Npr2p-Npr3p) subcomplex, a GAP for GTPase Gtr1p (EGOC subunit) in response to amino acid limitation, thereby resulting in activation of TORC1 signaling; SEA is a coatomer-related complex that associates dynamically with the vacuole; contains an N-terminal beta-propeller fold and a C-terminal RING motif |
| YBR093C | PHO5 | Repressible acid phosphatase |
| YBR181C | RPS6B | Protein component of the small (40S) ribosomal subunit; homologous to mammalian ribosomal protein S6, no bacterial homolog; phosphorylated on S233 by Ypk3p in a TORC1-dependent manner, and on S232 in a TORC1/2-dependent manner by Ypk1/2/3p; RPS6B has a paralog, RPS6A, that arose from the whole genome duplication |
| YBR266C | SLM6 | Protein with potential role in actin cytoskeleton organization; null mutant is slow-growing, sensitive to Congo Red, impaired in endocytosis, has increased lifespan; gene exhibits synthetic genetic interaction with MSS4 encoding phosphatidylinositol 4-phosphate kinase |
| YBR267W | REI1 | Cytoplasmic pre-60S factor; required for the correct recycling of shuttling factors Alb1, Arx1 and Tif6 at the end of the ribosomal large subunit biogenesis; involved in bud growth in the mitotic signaling network |
| YCL016C | DCC1 | Subunit of a complex with Ctf8p and Ctf18p |
| YCL064C | CHA1 | Catabolic L-serine (L-threonine) deaminase; catalyzes the degradation of both L-serine and L-threonine; required to use serine or threonine as the sole nitrogen source, transcriptionally induced by serine and threonine |
| YCR002C | CDC10 | Component of the septin ring, required for cytokinesis; septins are GTP-binding proteins that assemble into rod-like hetero-oligomers that can associate to form filaments; septin rings at the mother-bud neck act as scaffolds for recruiting cell division factors and as barriers to prevent diffusion of specific proteins between mother and daughter cells; N-terminus interacts with phosphatidylinositol-4,5-bisphosphate; protein abundance increases under DNA damage stress |
| YHR081W | LRP1 | Nuclear exosome-associated nucleic acid binding protein; involved in RNA processing, surveillance, degradation, tethering, and export; forms a stable heterodimer with Rrp6p and regulates its exonucleolytic activity; rapidly degraded by the proteasome in the absence of Rrp6p; homolog of mammalian nuclear matrix protein C1D involved in regulation of DNA repair and recombination |
| YHR120W | MSH1 | DNA-binding protein of the mitochondria |
| YKL048C | ELM1 | Serine/threonine protein kinase; regulates the orientation checkpoint, the morphogenesis checkpoint and the metabolic switch from fermentative to oxidative metabolism by phosphorylating the activation loop of Kin4p, Hsl1p and Snf1p respectively; cooperates with Hsl7p in recruiting Hsl1p to the septin ring, a prerequisite for subsequent recruitment, phosphorylation, and degradation of Swe1p; forms part of the bud neck ring; regulates cytokinesis |
| YKL110C | KTI12 | Protein that plays a role in modification of tRNA wobble nucleosides |
| YKL120W | OAC1 | Mitochondrial inner membrane transporter; transports oxaloacetate, sulfate, thiosulfate, and isopropylmalate; member of the mitochondrial carrier family |
| YKL191W | DPH2 | Protein required for synthesis of diphthamide; required along with Dph1p, Kti11p, Jjj3p, and Dph5p; diphthamide is a modified histidine residue of translation elongation factor 2 (Eft1p or Eft2p); forms a complex with Dph1p that catalyzes the first step of diphthamide biosynthesis |
| YKL213C | DOA1 | WD-repeat protein involved in Ub-mediated proteolysis; K48-, K29-specific Ub chain binding protein involved in Ub homeostasis; substrate-recruiting adaptor for Cdc48p in mitochondria-associated degradation; inhibits degradation of Ufd2p-dependent substrates; facilitates proteolysis of Cse4p, a centromeric H3-like protein; required for ribophagy; promotes NHEJ in postdiauxic/stationary phase; protein increases in abundance and relocates from nucleus to nuclear periphery upon replication stress |
| YKR035C | OPI8 | Dubious open reading frame; unlikely to encode a functional protein, based on available experimental and comparative sequence data; partially overlaps verified gene DID2/YKR035W-A |
| YKR047W |  | Dubious open reading frame; unlikely to encode a functional protein, based on available experimental and comparative sequence data; partially overlaps the verified gene NAP1 |
| YKR048C | NAP1 | Histone chaperone and assembly chaperone for Rps6a/b; involved in histone exchange, replacing histone H2A-H2B dimers or histone variant dimers from assembled nucleosomes; promotes solubility of Rps6a, and 40S ribosomal subunit biogenesis; role in transport of H2A and H2B histones to the nucleus; regulates microtubule dynamics during mitosis; interacts with mitotic cyclin Clb2p; controls bud morphogenesis; phosphorylated by CK2; protein abundance increases in response to DNA replication stress |
| YDL118W |  | Dubious open reading frame, unlikely to encode a protein; overlaps almost completely with YDL119C; mutations that would affect both YDL118W and YDL119C confer defective telomere maintenance and are synthetically sick or lethal with alpha-synuclein |
| YDR024W | FYV1 | Dubious open reading frame; unlikely to encode a functional protein, based on available experimental and comparative sequence data; not conserved in closely related Saccharomyces species; mutation decreases survival upon exposure to K1 killer toxin |
| YDR049W | VMS1 | Peptidyl-tRNA hydrolase that releases stalled peptides from ribosomes, component of a Cdc48p-complex; involved in protein quality control; exhibits cytosolic and ER-membrane localization; contributes to ER-associated degradation (ERAD) of specific substrates at a step after their ubiquitination; translocates to mitochondria under oxidative stress and forms a complex with Cdc48p and Npl4p that is required for ubiquitin-mediated mitochondria-associated protein degradation (MAD) |
| YDR080W | VPS41 | Subunit of the HOPS endocytic tethering complex; vacuole membrane protein that functions as a Rab GTPase effector, interacting specifically with the GTP-bound conformation of Ypt7p, facilitating tethering, docking and promoting membrane fusion events at the late endosome and vacuole; required for both membrane and protein trafficking; Yck3p-mediated phosphorylation regulates the organization of vacuolar fusion sites |
| YDR083W | RRP8 | Nucleolar S-adenosylmethionine-dependent rRNA methyltransferase |
| YDR127W | ARO1 | Pentafunctional arom protein; catalyzes steps 2 through 6 in the biosynthesis of chorismate, which is a precursor to aromatic amino acids |
| YDR128W | MTC5 | Subunit of SEACAT, a subcomplex of the SEA complex; Mtc1p, along with Rtc1p and Sea4p, redundantly inhibit the TORC1 inhibitory role of the Iml1p/SEACIT (Iml1p-Npr2p-Npr3p) subcomplex, a GAP for GTPase Gtr1p (EGOC subunit) in response to amino acid limitation, thereby resulting in activation of TORC1 signaling; SEA is a coatomer-related complex that associates dynamically with the vacuole; relative distribution to the vacuolar membrane decreases upon DNA replication stress |
| YDR156W | RPA14 | RNA polymerase I subunit A14 |
| YGR036C | CAX4 | Dolichyl pyrophosphate (Dol-P-P) phosphatase; cleaves the anhydride linkage in Dol-P-P; involved in synthesis of Dol-P-P-linked oligosaccharide intermediate required for protein N-glycosylation; has a lumenally oriented active site in the ER; mutation affects vesicular transport, membrane and cell wall biogenesis and lipid homeostasis |
| YGR057C | LST7 | Subunit of the Lst4p-Lst7p GTPase activating protein complex for Gtr2p; stimulates the GTPase activity of Rag family GTPase Gtr2p, within the context of the Gtr1p-Gtr2p heterodimer, after amino acid stimulation; required for activation of TORC1 in response to amino acid stimulation; recruited to the vacuolar membrane during amino acid starvation and released from the membrane by TORC1; required for the transport of amino acid permease Gap1p from the Golgi to the cell surface |
| YJL204C | RCY1 | F-box protein involved in recycling endocytosed proteins |
| YLR074C | BUD20 | C2H2-type zinc finger protein required for ribosome assembly; shuttling factor which associates with pre-60S particles in the nucleus, accompanying them to the cytoplasm; cytoplasmic dissociation of Bud20p requires Drg1p; N-terminus harbors a nuclear localization signal (NLS) and a nuclear export signal (NES); cytoplasmic Bud20p is reimported by Kap123-dependent pathway; involved in bud-site selection; diploid mutants display a random budding pattern; similar to human ZNF593 |
| YLR087C | CSF1 | Protein with structural similarity to lipid transport protein Vps13p |
| YLR089C | ALT1 | Alanine transaminase (glutamic pyruvic transaminase); involved in alanine biosynthesis and catabolism; TOR1-independent role in determining chronological lifespan; expression is induced in the presence of alanine; repression is mediated by Nrg1p; ALT1 has a paralog, ALT2, that arose from the whole genome duplication; Alt2p is catalytically inactive |
| YLR098C | CHA4 | DNA binding transcriptional activator |
| YLR244C | MAP1 | Methionine aminopeptidase; catalyzes the cotranslational removal of N-terminal methionine from nascent polypeptides; function is partially redundant with that of Map2p; apo-Map1p is activated by the GTP-dependent zinc transferase Zng1p, especially during zinc starvation; interacts with Zng1p via an N-terminal Zn finger domain |
| YLR261C | VPS63 | Putative protein of unknown function |
| YLR262C | YPT6 | Rab family GTPase; required for endosome-to-Golgi, intra-Golgi retrograde, and retrograde Golgi-to-ER transport; temporarily at the Golgi, dissociating into the cytosol on arrival of the late Golgi GTPase Ypt32p; Golgi-localized form is GTP bound, while cytosolic form is GDP-bound; required for delivery of Atg9p to the phagophore assembly site during autophagy under heat stress, with Ypt6p for starvation induced autophagy and for the CVT pathway; homolog of mammalian Rab6 |
| YLR335W | NUP2 | Nucleoporin involved in nucleocytoplasmic transport |
| YLR342W | FKS1 | Catalytic subunit of 1,3-beta-D-glucan synthase; transfers glucose from UDP-glucose to growing chains of glucan via β-1,3-glycosidic linkages, translocating polymerized β-1,3-glucan to the extracellular space; role in cell wall synthesis and maintenance; functionally redundant with alternate catalytic subunit Gsc2p; activity regulated by Rho1p, a subunit of the synthase complex; localizes to sites of cell wall remodeling; contains a cellulose-synthase-like fold; antifungal drug target |
| YLR386W | VAC14 | Enzyme regulator; involved in synthesis of phosphatidylinositol 3,5-bisphosphate, in control of trafficking of some proteins to the vacuole lumen via the MVB, and in maintenance of vacuole size and acidity; binds negative (Fig4p) and positive (Fab1p) regulators of PtdIns(3,5)P(2) to control endolysosome function; similar to mammalian Vac14p |
| YLR388W | RPS29A | Protein component of the small (40S) ribosomal subunit; homologous to mammalian ribosomal protein S29 and bacterial S14; RPS29A has a paralog, RPS29B, that arose from the whole genome duplication |
| YLR402W |  | Dubious open reading frame; unlikely to encode a functional protein, based on available experimental and comparative sequence data |
| YML001W | YPT7 | Rab family GTPase; GTP-binding protein of the rab family; required for homotypic fusion event in vacuole inheritance, for endosome-endosome fusion; localizes to sites of contact between the vacuole and mitochondria (vCLAMPs); interacts with the cargo selection/retromer complex for retrograde sorting; similar to mammalian Rab7 |
| YML009W-B |  | Dubious open reading frame; unlikely to encode a functional protein, based on available experimental and comparative sequence data; deletion mutation confers an increase in Ty1 transposition |
| YML012C-A |  | Dubious open reading frame; unlikely to encode a functional protein, based on available experimental and comparative sequence data; partially overlaps the verified gene SEL1 |
| YML013W | UBX2 | Bridging factor involved in ER-associated protein degradation (ERAD) |
| YML026C | RPS18B | Protein component of the small (40S) ribosomal subunit; homologous to mammalian ribosomal protein S18 and bacterial S13; RPS18B has a paralog, RPS18A, that arose from the whole genome duplication; protein abundance increases in response to DNA replication stress |
| YMR205C | PFK2 | Beta subunit of heterooctameric phosphofructokinase; involved in glycolysis; indispensable for anaerobic growth; activated by fructose-2,6-bisphosphate and AMP; mutation inhibits glucose induction of cell cycle-related genes |
| YMR216C | SKY1 | SR protein kinase (SRPK) |
| YMR275C | BUL1 | Alpha-arrestin, component of the Rsp5p E3-ubiquitin ligase complex; ubiquitin-binding adaptor involved in down-regulation of general amino acid transporter Gap1p upon shift to preferred nitrogen source; involved in lactate transporter Jen1p endocytosis; BUL1 has a paralog, BUL2, that arose from the whole genome duplication |
| YNL016W | PUB1 | Poly (A)+ RNA-binding protein; abundant mRNP-component protein that binds cell wall mRNAs; required for stability of many mRNAs; component of glucose deprivation induced stress granules, involved in P-body-dependent granule assembly; implicated in regulation of translation; carries Q/N-rich domain at C-terminus, identified as candidate prion; ribosome-bound Pub1p modulates stop codon decoding during translation termination |
| YNL136W | EAF7 | Subunit of the NuA4 histone acetyltransferase complex |
| YNL162W | RPL42A | Ribosomal 60S subunit protein L42A; homologous to mammalian ribosomal protein L36A, no bacterial homolog; RPL42A has a paralog, RPL42B, that arose from the whole genome duplication |
| YNL198C |  | Dubious open reading frame; unlikely to encode a functional protein, based on available experimental and comparative sequence data |
| YNL199C | GCR2 | Transcriptional activator of genes involved in glycolysis |
| YNL215W | IES2 | Protein that associates with the INO80 chromatin remodeling complex |
| YNL226W |  | Dubious open reading frame; unlikely to encode a functional protein, based on available experimental and comparative sequence data; overlaps the verified gene JJJ1/YNL227C |
| YNL294C | RIM21 | pH sensor molecule, component of RIM101 pathway; positive regulator of cell size; has role in cell wall construction and alkaline pH response; is glycosylated and phosphorylated; interacts with Dfg16p and Rim9p to form a pH-sensing complex; localization to plasma membrane is dependent on Dfg16p and Rim9p; has similarity to *A. nidulans* PalH |
| YNL307C | MCK1 | Dual-specificity S/T and tyrosine protein kinase; spindle position checkpoint component that works in parallel with Kin4p to prevent mitotic exit network (MEN) activation by the Cdc14 early anaphase release (FEAR) network; roles in chromosome segregation, meiotic entry, genome stability, phosphorylation-dependent degradation of Rcn1p and Cdc6p, inhibition of protein kinase A, transcriptional regulation, inhibition of RNAPIII, calcium stress and inhibition of Clb2p-Cdc28p after nuclear division |
| YNR010W | CSE2 | Subunit of the RNA polymerase II mediator complex |
| YOR001W | RRP6 | Nuclear exosome exonuclease component; has 3'-5' exonuclease activity that is regulated by Lrp1p; involved in RNA processing, maturation, surveillance, degradation, tethering, and export; role in sn/snoRNAs precursor degradation; forms a stable heterodimer with Lrp1p; has similarity to *E. coli* RNase D and to human PM-Sc1 100 (EXOSC10); mutant displays reduced transcription elongation in the G-less-based |
| YOR006C | TSR3 | Protein required for 20S pre-rRNA processing |
| YOR106W | VAM3 | Syntaxin-like vacuolar t-SNARE |
| YOR309C |  | Dubious open reading frame; unlikely to encode a functional protein, based on available experimental and comparative sequence data; partially overlaps the verified gene NOP58 |
| YPL029W | SUV3 | ATP-dependent RNA helicase |
| YPL180W | TCO89 | Subunit of TORC1 (Tor1p or Tor2p-Kog1p-Lst8p-Tco89p) |
| YPL226W | NEW1 | Translation termination and ribosome biogenesis factor |
| YPR119W | CLB2 | B-type cyclin involved in cell cycle progression; also involved in response to DNA damage; activates Cdc28p to promote the transition from G2 to M phase; accumulates during G2 and M, then targeted via a destruction box motif for ubiquitin-mediated degradation by the proteasome; CLB2 has a paralog, CLB1, that arose from the whole genome duplication |

**Supplementary Table 2:** All significantly enriched phenotypes.

| Phenotype | Outliers | Frequency | FC frequency | Z score | P-val. | adj.pval |
| --- | --- | --- | --- | --- | --- | --- |
| Media: minimal medium decreased competitive fitness | 53 | 0.779 | 0.468 | 3.618 | 1.48E-04 | 1.78E-03 |
| alpha-amino acid abnormal chemical compound accumulation | 50 | 0.735 | 1.353 | 8.517 | 8.18E-18 | 7.53E-16 |
| Media: YPD decreased competitive fitness | 48 | 0.706 | 1.524 | 8.574 | 4.98E-18 | 4.72E-16 |
| Media: ETH rich medium (YPD) w/6% ethanol decreased competitive fitness | 47 | 0.691 | 1.269 | 9.032 | 8.41E-20 | 8.77E-18 |
| Media: SC synthetic complete medium decreased competitive fitness | 45 | 0.662 | 1.152 | 6.385 | 8.59E-11 | 3.63E-09 |
| Media: OAK synthetic oak exudate medium decreased competitive fitness | 44 | 0.647 | 1.189 | 6.647 | 1.50E-11 | 6.80E-10 |
| decreased rate vegetative growth | 39 | 0.574 | 1.433 | 7.053 | 8.77E-13 | 5.83E-11 |
| Temperature: elevated temperature (37°C) increased heat sensitivity | 38 | 0.559 | 1.708 | 9.091 | 4.89E-20 | 5.27E-18 |
| Media: YPG glycerol medium decreased competitive fitness | 33 | 0.485 | 0.913 | 4.905 | 4.68E-07 | 9.72E-06 |
| ethanol decreased competitive fitness | 29 | 0.426 | 1.053 | 4.833 | 6.73E-07 | 1.33E-05 |
| histone deacetylase inhibitor CG-1521 7-phenyl-2,4,6-heptatrienoic hydroxamic acid decreased resistance to chemicals | 28 | 0.412 | 1.276 | 5.553 | 1.41E-08 | 3.67E-07 |
| alanine increased chemical compound accumulation | 26 | 0.382 | 2.434 | 10.298 | 3.59E-25 | 1.60E-22 |
| Media: 16 h YPD decreased competitive fitness | 26 | 0.382 | 2.154 | 9.751 | 9.13E-23 | 1.14E-20 |
| 1 uM zinc dichloride Media: zinc deficiency increased stress resistance | 26 | 0.382 | 2.225 | 9.715 | 1.30E-22 | 1.56E-20 |
| decreased chronological lifespan | 24 | 0.353 | 0.804 | 2.991 | 1.39E-03 | 1.26E-02 |
| 460 uM dieldrin increased resistance to chemicals | 21 | 0.309 | 2.325 | 8.288 | 5.77E-17 | 5.01E-15 |
| Temperature: elevated temperature decreased cell death | 21 | 0.309 | 0.805 | 3.014 | 1.29E-03 | 1.19E-02 |
| 50 ug/ml Congo Red decreased resistance to chemicals | 20 | 0.294 | 1.819 | 6.32 | 1.31E-10 | 5.31E-09 |
| abnormal protein/peptide distribution | 20 | 0.294 | 1.569 | 5.865 | 2.25E-09 | 7.74E-08 |
| increased chronological lifespan | 19 | 0.279 | 1.147 | 3.824 | 6.56E-05 | 8.51E-04 |
| Media: YPD decreased rate fermentative growth | 18 | 0.265 | 2.434 | 8.497 | 9.77E-18 | 8.73E-16 |
| 10 uM benomyl increased resistance to chemicals | 18 | 0.265 | 1.99 | 6.957 | 1.74E-12 | 8.38E-11 |
| 1 mM paraquat decreased oxidative stress resistance | 18 | 0.265 | 1.404 | 4.733 | 1.11E-06 | 2.10E-05 |
| 3 mM hydrogen peroxide increased oxidative stress resistance | 17 | 0.25 | 1.861 | 6.704 | 1.02E-11 | 4.75E-10 |
| Treatment: pooled mutants subject to three cycles of growth to post-diauxic phase followed by desiccation and regrowth decreased desiccation resistance | 17 | 0.25 | 0.986 | 3.262 | 5.54E-04 | 5.73E-03 |
| 3 ug/ml Calcofluor White decreased resistance to chemicals | 16 | 0.235 | 2.672 | 8.876 | 3.47E-19 | 3.39E-17 |
| 50 ug/ml hygromycin B decreased resistance to chemicals | 16 | 0.235 | 2.317 | 7.503 | 3.12E-14 | 2.27E-12 |
| myo-inositol auxotrophy | 16 | 0.235 | 1.522 | 4.69 | 1.36E-06 | 2.54E-05 |
| 2 mg/ml caffeine decreased resistance to chemicals | 15 | 0.221 | 2.809 | 9.28 | 8.48E-21 | 9.47E-19 |
| decreased cell size | 15 | 0.221 | 2.265 | 6.291 | 1.57E-10 | 6.30E-09 |
| 1 uM sodium selenide decreased resistance to chemicals | 15 | 0.221 | 2.124 | 5.988 | 1.06E-09 | 3.94E-08 |
| decreased transposable element transposition | 15 | 0.221 | 1.903 | 5.849 | 2.47E-09 | 8.38E-08 |
| sirolimus decreased resistance to chemicals | 15 | 0.221 | 1.893 | 5.624 | 9.35E-09 | 2.48E-07 |
| 8% ethanol decreased resistance to chemicals | 15 | 0.221 | 0.868 | 2.72 | 3.26E-03 | 2.62E-02 |
| polyphosphate abnormal chemical compound accumulation | 14 | 0.206 | 2.055 | 5.894 | 1.89E-09 | 6.56E-08 |
| decreased endocytosis | 14 | 0.206 | 1.432 | 4.032 | 2.76E-05 | 3.89E-04 |
| 30% sucrose decreased hyperosmotic stress resistance | 13 | 0.191 | 2.145 | 6.267 | 1.83E-10 | 7.17E-09 |
| 7.7 mM caffeine decreased resistance to chemicals | 13 | 0.191 | 1.69 | 4.572 | 2.41E-06 | 4.34E-05 |
| 100 mM hydroxyurea decreased resistance to chemicals | 13 | 0.191 | 1.393 | 3.87 | 5.45E-05 | 7.31E-04 |
| spermine decreased resistance to chemicals | 12 | 0.176 | 2.508 | 7.104 | 6.04E-13 | 4.20E-11 |
| decreased killer toxin resistance | 12 | 0.176 | 2.481 | 7.022 | 1.09E-12 | 6.25E-11 |
| hygromycin B decreased resistance to chemicals | 12 | 0.176 | 2.447 | 6.577 | 2.39E-11 | 1.05E-09 |
| 489.5 µM N-nitrosodiethylamine decreased resistance to chemicals | 12 | 0.176 | 1.939 | 5.758 | 4.26E-09 | 1.37E-07 |
| decreased replicative lifespan | 12 | 0.176 | 1.585 | 4.634 | 1.79E-06 | 3.28E-05 |
| 0.015 µg/ml sirolimus decreased resistance to chemicals | 12 | 0.176 | 1.925 | 4.503 | 3.35E-06 | 5.86E-05 |
| Treatment: 400-700nm visible light decreased stress resistance | 12 | 0.176 | 0.939 | 2.5 | 6.21E-03 | 4.63E-02 |
| 15 µg/ml 5-fluorouracil decreased resistance to chemicals | 11 | 0.162 | 2.964 | 7.878 | 1.66E-15 | 1.30E-13 |
| increased duration cell cycle progression in G1 phase | 11 | 0.162 | 2.362 | 6.328 | 1.24E-10 | 5.11E-09 |
| 0.4% (w/v), pH 4.3 acetic acid decreased resistance to chemicals | 11 | 0.162 | 2.309 | 6.221 | 2.46E-10 | 9.51E-09 |
| glutamate increased chemical compound accumulation | 11 | 0.162 | 2.173 | 6.2 | 2.83E-10 | 1.08E-08 |
| 12 mM sodium disulfite decreased resistance to chemicals | 11 | 0.162 | 2.29 | 5.735 | 4.86E-09 | 1.55E-07 |
| 1772 µM methyl methanesulfonate decreased resistance to chemicals | 11 | 0.162 | 2.059 | 5.097 | 1.73E-07 | 4.03E-06 |
| 3.75 mM yttrium chloride increased resistance to chemicals | 11 | 0.162 | 1.715 | 4.566 | 2.49E-06 | 4.42E-05 |
| 16.5 µM fenpropimorph increased resistance to chemicals | 11 | 0.162 | 1.669 | 4.515 | 3.16E-06 | 5.55E-05 |
| 7 µg/ml hygromycin B decreased resistance to chemicals | 11 | 0.162 | 1.841 | 4.472 | 3.88E-06 | 6.67E-05 |
| decreased protein/peptide accumulation | 11 | 0.162 | 1.681 | 4.396 | 5.51E-06 | 9.26E-05 |
| 40 µM cadmium(2+) decreased metal resistance | 11 | 0.162 | 1.795 | 4.284 | 9.19E-06 | 1.37E-04 |
| 50 µM idarubicin increased resistance to chemicals | 11 | 0.162 | 1.56 | 4.213 | 1.26E-05 | 1.85E-04 |
| 16.4 mM 4-nitroquinoline N-oxide decreased resistance to chemicals | 11 | 0.162 | 1.576 | 3.845 | 6.02E-05 | 7.95E-04 |
| Temperature: elevated temperature (39°C) increased heat sensitivity | 11 | 0.162 | 1.624 | 3.632 | 1.41E-04 | 1.71E-03 |
| glutamate(1-) decreased rate utilization of nitrogen source | 11 | 0.162 | 1.264 | 3.301 | 4.83E-04 | 5.08E-03 |
| decreased desiccation resistance | 11 | 0.162 | 1.264 | 3.109 | 9.38E-04 | 8.92E-03 |
| 46.8 µM mycophenolic acid increased resistance to chemicals | 10 | 0.147 | 2.184 | 5.782 | 3.70E-09 | 1.20E-07 |
| 2 µg/ml bleomycin A5 decreased resistance to chemicals | 10 | 0.147 | 2.198 | 5.603 | 1.05E-08 | 2.76E-07 |
| 1.5 g/L quinine decreased resistance to chemicals | 10 | 0.147 | 1.781 | 5.282 | 6.40E-08 | 1.57E-06 |
| 250 µM tirapazamine increased resistance to chemicals | 10 | 0.147 | 1.878 | 4.884 | 5.19E-07 | 1.04E-05 |
| 400 µM actinomycin D decreased resistance to chemicals | 10 | 0.147 | 1.766 | 4.853 | 6.08E-07 | 1.21E-05 |
| hydroxyurea decreased resistance to chemicals | 10 | 0.147 | 1.9 | 4.524 | 3.03E-06 | 5.35E-05 |
| 0.6 µM tunicamycin decreased resistance to chemicals | 10 | 0.147 | 1.989 | 4.385 | 5.80E-06 | 9.64E-05 |
| Treatment: lignocellulosic wheat straw hydrolysate (WSH) decreased vegetative growth | 10 | 0.147 | 1.617 | 4.322 | 7.75E-06 | 1.16E-04 |
| 50 µg/ml doxorubicin decreased resistance to chemicals | 10 | 0.147 | 1.766 | 4.134 | 1.78E-05 | 2.58E-04 |
| 200 µg/ml sulforaphane decreased resistance to chemicals | 10 | 0.147 | 1.718 | 3.967 | 3.64E-05 | 5.06E-04 |
| 0.03% methyl methanesulfonate increased resistance to chemicals | 10 | 0.147 | 1.626 | 3.813 | 6.85E-05 | 8.85E-04 |
| 3.2 µM bleomycin increased resistance to chemicals | 10 | 0.147 | 1.381 | 3.13 | 8.75E-04 | 8.37E-03 |
| 20.53 µM penconazole increased resistance to chemicals | 9 | 0.132 | 3.244 | 10.004 | 7.31E-24 | 1.64E-21 |
| 444 mM formamide decreased resistance to chemicals | 9 | 0.132 | 3.259 | 8.157 | 1.72E-16 | 1.45E-14 |
| abnormal endoplasmic reticulum morphology | 9 | 0.132 | 2.388 | 5.941 | 1.42E-09 | 5.09E-08 |
| 6 mM caffeine decreased resistance to chemicals | 9 | 0.132 | 2.12 | 5.504 | 1.85E-08 | 4.75E-07 |
| Phase: stationary phase, Treatment: 50°C heat shock increased innate thermotolerance | 9 | 0.132 | 2.338 | 5.168 | 1.18E-07 | 2.84E-06 |
| serine decreased chemical compound accumulation | 9 | 0.132 | 2.221 | 5.122 | 1.51E-07 | 3.55E-06 |
| abnormal bud morphology | 9 | 0.132 | 2.046 | 4.938 | 3.95E-07 | 8.88E-06 |
| 0.15% caffeine decreased resistance to chemicals | 9 | 0.132 | 2.214 | 4.899 | 4.80E-07 | 9.72E-06 |
| 6.4 mM methylamine decreased resistance to chemicals | 9 | 0.132 | 1.925 | 3.77 | 8.17E-05 | 1.03E-03 |
| 100 µM fluconazole increased resistance to chemicals | 9 | 0.132 | 1.515 | 3.623 | 1.46E-04 | 1.76E-03 |
| 0.75 - 1.0 mM arsenite(3-) decreased resistance to chemicals | 9 | 0.132 | 1.469 | 3.507 | 2.26E-04 | 2.60E-03 |
| 43 µM camptothecin decreased resistance to chemicals | 9 | 0.132 | 1.33 | 3.226 | 6.29E-04 | 6.40E-03 |
| 0.4 mM aluminium(3+), 0.8 mM 1,1'-azobis(N,N-dimethylformamide) decreased resistance to chemicals | 9 | 0.132 | 1.496 | 3.192 | 7.07E-04 | 7.11E-03 |
| Treatment: Aβ42 decreased toxin resistance | 9 | 0.132 | 1.31 | 3.034 | 1.21E-03 | 1.13E-02 |
| 8% dimethyl sulfoxide decreased resistance to chemicals | 9 | 0.132 | 1.214 | 2.591 | 4.79E-03 | 3.68E-02 |
| chondramide derivative decreased resistance to chemicals | 9 | 0.132 | 1.145 | 2.575 | 5.01E-03 | 3.82E-02 |
| arginine increased chemical compound accumulation | 8 | 0.118 | 2.201 | 4.756 | 9.87E-07 | 1.90E-05 |
| increased cell size | 8 | 0.118 | 2.051 | 4.676 | 1.46E-06 | 2.69E-05 |
| 0.04 µg/mL flucytosine decreased resistance to chemicals | 8 | 0.118 | 1.979 | 4.344 | 7.01E-06 | 1.07E-04 |
| valine increased chemical compound accumulation | 8 | 0.118 | 1.805 | 3.827 | 6.48E-05 | 8.48E-04 |
| hydrogen sulfide Media: 500 mg/L cysteine supplementation decreased chemical compound accumulation | 8 | 0.118 | 1.599 | 3.605 | 1.56E-04 | 1.86E-03 |
| 50 μM bis(S-citronellalthiosemicarbazonato)nickel(II) decreased resistance to chemicals | 8 | 0.118 | 1.444 | 3.399 | 3.38E-04 | 3.77E-03 |
| 100 μM fluconazole decreased resistance to chemicals | 8 | 0.118 | 1.494 | 3.377 | 3.66E-04 | 3.94E-03 |
| 0.0375 mg/ml drimenol decreased resistance to chemicals | 8 | 0.118 | 1.474 | 3.261 | 5.56E-04 | 5.74E-03 |
| glutathione Treatment: SD medium increased chemical compound excretion | 8 | 0.118 | 1.336 | 2.928 | 1.71E-03 | 1.52E-02 |
| 0.6 μM tunicamycin increased resistance to chemicals | 8 | 0.118 | 1.43 | 2.612 | 4.50E-03 | 3.49E-02 |

**Supplementary Table 3:**
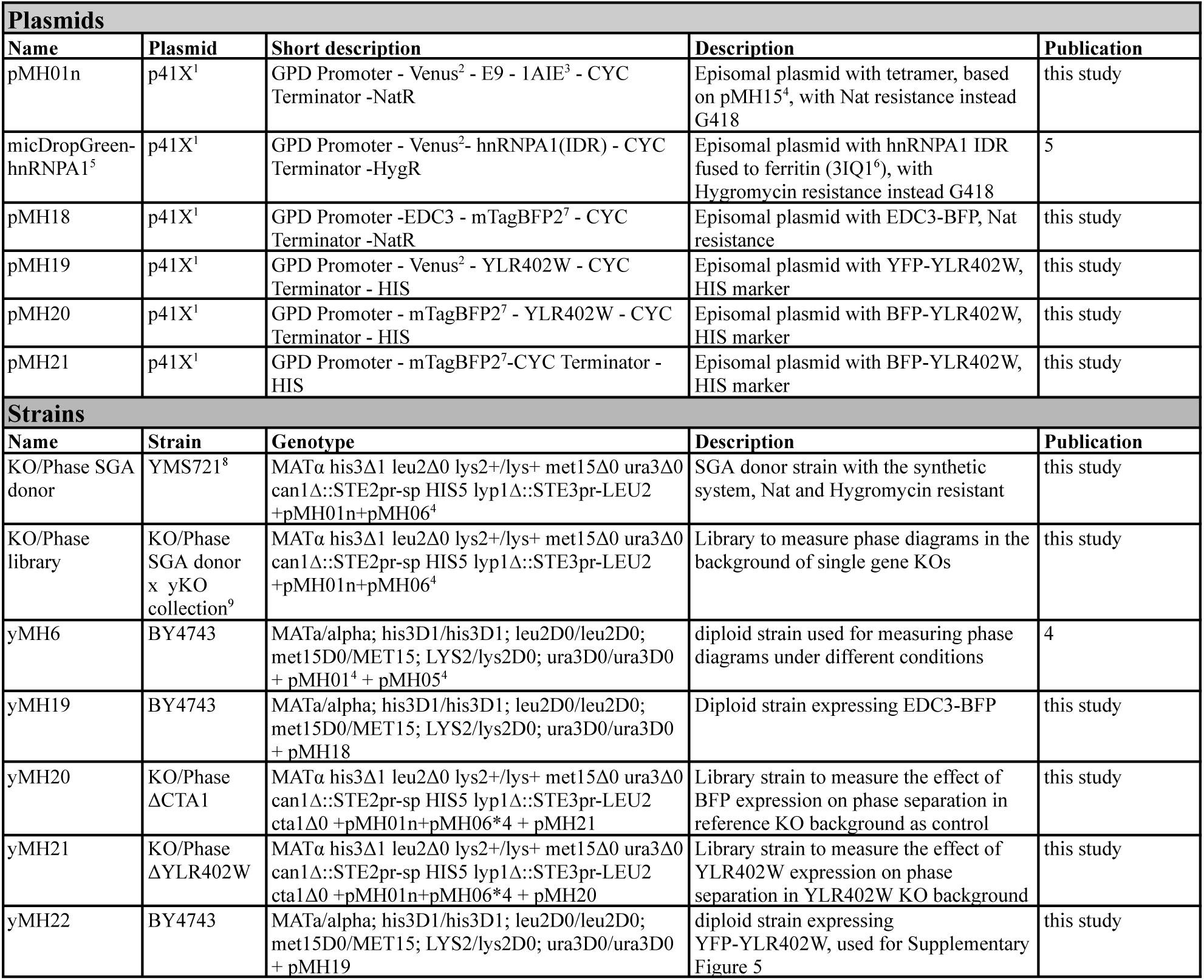
Plasmids and strains used and generated in this study. References: 1 (Mumberg, Muller, and Funk 1995), 2 (Nagai et al. 2002), 3 (Mittal et al. 1998), 4 (Heidenreich et al. 2020), 5 (Gilat et al. 2023), 6 (Nocek et al. 2009), 7 (Subach et al. 2011), 8 (Breslow et al. 2008), 9 (Wach et al. 1994; Winzeler 1999).

## Notes

https://figshare.com/s/1b2e878ef2e26f8b1c80

### Competing Interest Statement

The authors have declared no competing interest.

## References

1. S. Boeynaems, S. Alberti, N. L. Fawzi, T. Mittag, M. Polymenidou, F. Rousseau, J. Schymkowitz, J. Shorter, B. Wolozin, L. Van Den Bosch, P. Tompa, M. Fuxreiter, Protein Phase Separation: A New Phase in Cell Biology. Trends Cell Biol. 28, 420–435 (2018).

2. D. M. Mitrea, R. W. Kriwacki, Phase separation in biology; functional organization of a higher order. Cell Commun. Signal. 14, 1 (2016).

3. C. P. Brangwynne, C. R. Eckmann, D. S. Courson, A. Rybarska, C. Hoege, J. Gharakhani, F. Jülicher, A. A. Hyman, Germline P granules are liquid droplets that localize by controlled dissolution/condensation. Science 324, 1729–1732 (2009).

4. S. F. Banani, H. O. Lee, A. A. Hyman, M. K. Rosen, Biomolecular condensates: organizers of cellular biochemistry. Nat. Rev. Mol. Cell Biol. 18, 285–298 (2017).

5. B. Wang, L. Zhang, T. Dai, Z. Qin, H. Lu, L. Zhang, F. Zhou, Liquid–liquid phase separation in human health and diseases. Signal Transduction and Targeted Therapy 6, 1–16 (2021).

6. S. Alberti, D. Dormann, Liquid-Liquid Phase Separation in Disease. Annu. Rev. Genet. 53, 171–194 (2019).

7. A. Patel, L. Malinovska, S. Saha, J. Wang, S. Alberti, Y. Krishnan, A. A. Hyman, ATP as a biological hydrotrope. Science 356, 753–756 (2017).

8. J. A. Villegas, M. Heidenreich, E. D. Levy, Molecular and environmental determinants of biomolecular condensate formation. Nat. Chem. Biol. 18, 1319–1329 (2022).

9. M. Delarue, G. P. Brittingham, S. Pfeffer, I. V. Surovtsev, S. Pinglay, K. J. Kennedy, M. Schaffer, J. I. Gutierrez, D. Sang, G. Poterewicz, J. K. Chung, J. M. Plitzko, J. T. Groves, C. Jacobs-Wagner, B. D. Engel, L. J. Holt, mTORC1 Controls Phase Separation and the Biophysical Properties of the Cytoplasm by Tuning Crowding. Cell 174, 338–349.e20 (2018).

10. C. Garcia-Cabau, X. Salvatella, Regulation of biomolecular condensate dynamics by signaling. Curr. Opin. Cell Biol. 69, 111–119 (2021).

11. B. R. Parry, I. V. Surovtsev, M. T. Cabeen, C. S. O’Hern, E. R. Dufresne, C. Jacobs-Wagner, The bacterial cytoplasm has glass-like properties and is fluidized by metabolic activity. Cell 156, 183–194 (2014).

12. M. B. Heimlicher, M. Bächler, M. Liu, C. Ibeneche-Nnewihe, E.-L. Florin, A. Hoenger, D. Brunner, Reversible solidification of fission yeast cytoplasm after prolonged nutrient starvation. J. Cell Sci. 132 (2019).

13. M. C. Munder, D. Midtvedt, T. Franzmann, E. Nüske, O. Otto, M. Herbig, E. Ulbricht, P. Müller, A. Taubenberger, S. Maharana, L. Malinovska, D. Richter, J. Guck, V. Zaburdaev, S. Alberti, A pH-driven transition of the cytoplasm from a fluid-to a solid-like state promotes entry into dormancy. Elife 5 (2016).

14. R. P. Joyner, J. H. Tang, J. Helenius, E. Dultz, C. Brune, L. J. Holt, S. Huet, D. J. Müller, K. Weis, A glucose-starvation response regulates the diffusion of macromolecules. Elife 5 (2016).

15. R. Narayanaswamy, M. Levy, M. Tsechansky, G. M. Stovall, J. D. O’Connell, J. Mirrielees, A. D. Ellington, E. M. Marcotte, Widespread reorganization of metabolic enzymes into reversible assemblies upon nutrient starvation. Proc. Natl. Acad. Sci. U. S. A. 106, 10147–10152 (2009).

16. D. Laporte, B. Salin, B. Daignan-Fornier, I. Sagot, Reversible cytoplasmic localization of the proteasome in quiescent yeast cells. J. Cell Biol. 181, 737–745 (2008).

17. I. Petrovska, E. Nüske, M. C. Munder, G. Kulasegaran, L. Malinovska, S. Kroschwald, D. Richter, K. Fahmy, K. Gibson, J.-M. Verbavatz, S. Alberti, Filament formation by metabolic enzymes is a specific adaptation to an advanced state of cellular starvation. [Preprint] (2014). 10.7554/elife.02409.

18. G. Marini, E. Nüske, W. Leng, S. Alberti, G. Pigino, Reorganization of budding yeast cytoplasm upon energy depletion. Mol. Biol. Cell 31, 1232–1245 (2020).

19. M. C. Munder, D. Midtvedt, T. Franzmann, E. Nüske, O. Otto, M. Herbig, E. Ulbricht, P. Müller, A. Taubenberger, S. Maharana, L. Malinovska, D. Richter, J. Guck, V. Zaburdaev, S. Alberti, A pH-driven transition of the cytoplasm from a fluid-to a solid-like state promotes entry into dormancy. Elife 5 (2016).

20. Y. Xiang, I. V. Surovtsev, Y. Chang, S. K. Govers, B. R. Parry, J. Liu, C. Jacobs-Wagner, Interconnecting solvent quality, transcription, and chromosome folding in Escherichia coli. Cell 184, 3626–3642.e14 (2021).

21. M. Delarue, G. P. Brittingham, S. Pfeffer, I. V. Surovtsev, S. Pinglay, K. J. Kennedy, M. Schaffer, J. I. Gutierrez, D. Sang, G. Poterewicz, J. K. Chung, J. M. Plitzko, J. T. Groves, C. Jacobs-Wagner, B. D. Engel, L. J. Holt, mTORC1 Controls Phase Separation and the Biophysical Properties of the Cytoplasm by Tuning Crowding. Cell 174, 338–349.e20 (2018).

22. A. Gilat, B. Dubrueil, E. D. Levy, Mapping interactions between disordered regions reveals promiscuity in biomolecular condensate formation, bioRxiv (2023)p. 2023.07.04.547715.

23. M. Heidenreich, J. M. Georgeson, E. Locatelli, L. Rovigatti, S. K. Nandi, A. Steinberg, Y. Nadav, E. Shimoni, S. A. Safran, J. P. K. Doye, E. D. Levy, Designer protein assemblies with tunable phase diagrams in living cells. Nat. Chem. Biol. 16, 939–945 (2020).

24. R. Orij, J. Postmus, A. Ter Beek, S. Brul, G. J. Smits, In vivo measurement of cytosolic and mitochondrial pH using a pH-sensitive GFP derivative in Saccharomyces cerevisiae reveals a relation between intracellular pH and growth. Microbiology 155, 268–278 (2009).

25. Y. Xie, T. Shu, T. Liu, M.-C. Spindler, J. Mahamid, G. M. Hocky, D. Gresham, L. J. Holt, Polysome collapse and RNA condensation fluidize the cytoplasm. Mol. Cell 84, 2698–2716.e9 (2024).

26. R. Dechant, M. Binda, S. S. Lee, S. Pelet, J. Winderickx, M. Peter, Cytosolic pH is a second messenger for glucose and regulates the PKA pathway through V-ATPase. EMBO J. 29, 2515–2526 (2010).

27. A. H. Y. Tong, C. Boone, Synthetic genetic array analysis in Saccharomyces cerevisiae. Methods Mol. Biol. 313, 171–192 (2006).

28. E. A. Winzeler, Functional Characterization of the S. cerevisiae Genome by Gene Deletion and Parallel Analysis. [Preprint] (1999). 10.1126/science.285.5429.901.

29. Y. Cohen, M. Schuldiner, “Advanced Methods for High-Throughput Microscopy Screening of Genetically Modified Yeast Libraries” in Network Biology: Methods and Applications, G. Cagney, A. Emili, Eds. (Humana Press, Totowa, NJ, 2011), pp. 127–159.

30. J. Peter, M. De Chiara, A. Friedrich, J.-X. Yue, D. Pflieger, A. Bergström, A. Sigwalt, B. Barre, K. Freel, A. Llored, C. Cruaud, K. Labadie, J.-M. Aury, B. Istace, K. Lebrigand, P. Barbry, S. Engelen, A. Lemainque, P. Wincker, G. Liti, J. Schacherer, Genome evolution across 1,011 Saccharomyces cerevisiae isolates. Nature 556, 339–344 (2018).

31. M. Ashburner, C. A. Ball, J. A. Blake, D. Botstein, H. Butler, J. M. Cherry, A. P. Davis, K. Dolinski, S. S. Dwight, J. T. Eppig, M. A. Harris, D. P. Hill, L. Issel-Tarver, A. Kasarskis, S. Lewis, J. C. Matese, J. E. Richardson, M. Ringwald, G. M. Rubin, G. Sherlock, Gene ontology: tool for the unification of biology. The Gene Ontology Consortium. Nat. Genet. 25, 25–29 (2000).

32. Gene Ontology Consortium, The Gene Ontology resource: enriching a GOld mine. Nucleic Acids Res. 49, D325–D334 (2021).

33. R. Balakrishnan, J. Park, K. Karra, B. C. Hitz, G. Binkley, E. L. Hong, J. Sullivan, G. Micklem, J. Michael Cherry, YeastMine—an integrated data warehouse for Saccharomyces cerevisiae data as a multipurpose tool-kit. [Preprint] (2012). 10.1093/database/bar062.

34. Y. Luo, Z. Na, S. A. Slavoff, P-Bodies: Composition, Properties, and Functions. Biochemistry 57, 2424–2431 (2018).

35. D. Kaganovich, R. Kopito, J. Frydman, Misfolded proteins partition between two distinct quality control compartments. Nature 454, 1088–1095 (2008).

36. U. Raudvere, L. Kolberg, I. Kuzmin, T. Arak, P. Adler, H. Peterson, J. Vilo, g:Profiler: a web server for functional enrichment analysis and conversions of gene lists (2019 update). Nucleic Acids Res. 47, W191–W198 (2019).

37. L. B. Persson, V. S. Ambati, O. Brandman, Cellular Control of Viscosity Counters Changes in Temperature and Energy Availability. Cell 183 (2020).

38. C. B. Messner, V. Demichev, J. Muenzner, S. K. Aulakh, N. Barthel, A. Röhl, L. Herrera-Domínguez, A.-S. Egger, S. Kamrad, J. Hou, G. Tan, O. Lemke, E. Calvani, L. Szyrwiel, M. Mülleder, K. S. Lilley, C. Boone, G. Kustatscher, M. Ralser, The proteomic landscape of genome-wide genetic perturbations. Cell 186, 2018–2034.e21 (2023).

39. A. Agrawal, H. Balcı, K. Hanspers, S. L. Coort, M. Martens, D. N. Slenter, F. Ehrhart, D. Digles, A. Waagmeester, I. Wassink, T. Abbassi-Daloii, E. N. Lopes, A. Iyer, J. M. Acosta, L. G. Willighagen, K. Nishida, A. Riutta, H. Basaric, C. T. Evelo, E. L. Willighagen, M. Kutmon, A. R. Pico, WikiPathways 2024: next generation pathway database. Nucleic Acids Res. 52, D679–D689 (2024).

40. M. Kanehisa, M. Furumichi, Y. Sato, M. Kawashima, M. Ishiguro-Watanabe, KEGG for taxonomy-based analysis of pathways and genomes. Nucleic Acids Res. 51, D587–D592 (2023).

41. J. I. Gutierrez, G. P. Brittingham, Y. Karadeniz, K. D. Tran, A. Dutta, A. S. Holehouse, C. L. Peterson, L. J. Holt, SWI/SNF senses carbon starvation with a pH-sensitive low-complexity sequence. Elife 11 (2022).

42. I. Sagot, D. Laporte, The cell biology of quiescent yeast - a diversity of individual scenarios. J. Cell Sci. 132 (2019).

43. A. Wach, A. Brachat, R. Pöhlmann, P. Philippsen, New heterologous modules for classical or PCR-based gene disruptions in Saccharomyces cerevisiae. Yeast 10, 1793–1808 (1994).

44. H. Lyons, R. T. Veettil, P. Pradhan, C. Fornero, N. De La Cruz, K. Ito, M. Eppert, R. G. Roeder, B. R. Sabari, Functional partitioning of transcriptional regulators by patterned charge blocks. Cell 186, 327–345.e28 (2023).

45. X. Su, J. A. Ditlev, E. Hui, W. Xing, S. Banjade, J. Okrut, D. S. King, J. Taunton, M. K. Rosen, R. D. Vale, Phase separation of signaling molecules promotes T cell receptor signal transduction. Science 352, 595–599 (2016).

46. P. Li, S. Banjade, H.-C. Cheng, S. Kim, B. Chen, L. Guo, M. Llaguno, J. V. Hollingsworth, D. S. King, S. F. Banani, P. S. Russo, Q.-X. Jiang, B. T. Nixon, M. K. Rosen, Phase transitions in the assembly of multivalent signalling proteins. Nature 483, 336–340 (2012).

47. L. B. Case, J. A. Ditlev, M. K. Rosen, Regulation of Transmembrane Signaling by Phase Separation. Annu. Rev. Biophys. 48, 465–494 (2019).

48. A. M. Küffner, M. Prodan, R. Zuccarini, U. C. Palmiero, L. Faltova, P. Arosio, Acceleration of an Enzymatic Reaction in Liquid Phase Separated Compartments Based on Intrinsically Disordered Protein Domains. ChemSystemsChem 2, e2000001 (2020).

49. J. Jeong, J. H. Lee, C. C. Carcamo, M. W. Parker, J. M. Berger, DNA-Stimulated Liquid-Liquid phase separation by eukaryotic topoisomerase ii modulates catalytic function. Elife 11 (2022).

50. W. Peeples, M. K. Rosen, Mechanistic dissection of increased enzymatic rate in a phase-separated compartment. Nat. Chem. Biol. 17, 693–702 (2021).

51. J. A. Riback, C. D. Katanski, J. L. Kear-Scott, E. V. Pilipenko, A. E. Rojek, T. R. Sosnick, D. A. Drummond, Stress-Triggered Phase Separation Is an Adaptive, Evolutionarily Tuned Response. Cell 168, 1028–1040.e19 (2017).

52. D. T. McSwiggen, M. Mir, X. Darzacq, R. Tjian, Evaluating phase separation in live cells: diagnosis, caveats, and functional consequences. Genes Dev. 33, 1619–1634 (2019).

53. F. Pessina, U. Gioia, O. Brandi, S. Farina, M. Ceccon, S. Francia, F. d’Adda di Fagagna, DNA Damage Triggers a New Phase in Neurodegeneration. Trends Genet. 37, 337–354 (2021).

54. T. Mittag, R. V. Pappu, A conceptual framework for understanding phase separation and addressing open questions and challenges. Mol. Cell 82, 2201–2214 (2022).

55. A. M. Miangolarra, A. Duperray-Susini, M. Coppey, M. Castellana, Two timescales control the creation of large protein aggregates in cells. Biophys. J. 120, 2394–2399 (2021).

56. T. Shu, G. Mitra, J. B. Alberts, M. Viana, E. D. Levy, G. M. Hocky, L. J. Holt, Mesoscale molecular assembly is favored by the active, crowded cytoplasm. American Physical Society, doi: 10.1103/PRXLife.2.033001 (2024).

57. Y. Xie, T. Liu, D. Gresham, L. J. Holt, mRNA condensation fluidizes the cytoplasm. *bioRxiv*, doi: 10.1101/2023.05.30.542963 (2023).

58. J. D. Wurtz, C. F. Lee, Stress granule formation via ATP depletion-triggered phase separation. New J. Phys. 20, 045008 (2018).

59. M. Takaine, H. Imamura, S. Yoshida, High and stable ATP levels prevent aberrant intracellular protein aggregation in yeast. Elife 11 (2022).

60. J. L. Watson, E. Seinkmane, C. T. Styles, A. Mihut, L. K. Krüger, K. E. McNally, V. J. Planelles-Herrero, M. Dudek, P. M. McCall, S. Barbiero, M. Vanden Oever, S. Y. Peak-Chew, B. T. Porebski, A. Zeng, N. M. Rzechorzek, D. C. S. Wong, A. D. Beale, A. Stangherlin, M. Riggi, J. Iwasa, J. Morf, C. Miliotis, A. Guna, A. J. Inglis, J. Brugués, R. M. Voorhees, J. E. Chambers, Q.-J. Meng, J. S. O’Neill, R. S. Edgar, E. Derivery, Macromolecular condensation buffers intracellular water potential. Nature 623, 842–852 (2023).

61. J. A. Villegas, E. D. Levy, A unified statistical potential reveals that amino acid stickiness governs nonspecific recruitment of client proteins into condensates. Protein Sci. 31, e4361 (2022).

62. H. Roman, A system selective for mutations affecting the synthesis of adenine in yeast. Cr. Trav. Lab. Carlsberg Ser. Physiol. (1956).

63. S. Xie, M. Swaffer, J. M. Skotheim, Eukaryotic Cell Size Control and Its Relation to Biosynthesis and Senescence. Annu. Rev. Cell Dev. Biol. 38, 291–319 (2022).

64. L. H. Hartwell, J. J. Hopfield, S. Leibler, A. W. Murray, From molecular to modular cell biology. Nature 402, C47–52 (1999).

65. M. Costanzo, A. Baryshnikova, J. Bellay, Y. Kim, E. D. Spear, C. S. Sevier, H. Ding, J. L. Koh, K. Toufighi, S. Mostafavi, J. Prinz, R. P. St Onge, B. VanderSluis, T. Makhnevych, F. J. Vizeacoumar, S. Alizadeh, S. Bahr, R. L. Brost, Y. Chen, M. Cokol, R. Deshpande, Z. Li, Z. Y. Lin, W. Liang, M. Marback, J. Paw, B. J. San Luis, E. Shuteriqi, A. H. Tong, N. van Dyk, I. M. Wallace, J. A. Whitney, M. T. Weirauch, G. Zhong, H. Zhu, W. A. Houry, M. Brudno, S. Ragibizadeh, B. Papp, C. Pal, F. P. Roth, G. Giaever, C. Nislow, O. G. Troyanskaya, H. Bussey, G. D. Bader, A. C. Gingras, Q. D. Morris, P. M. Kim, C. A. Kaiser, C. L. Myers, B. J. Andrews, C. Boone, The genetic landscape of a cell. Science 327, 425–431 (2010).

66. Y. T. Chong, J. L. Koh, H. Friesen, S. K. Duffy, M. J. Cox, A. Moses, J. Moffat, C. Boone, B. J. Andrews, Yeast Proteome Dynamics from Single Cell Imaging and Automated Analysis. Cell 161, 1413–1424 (2015).

67. M. Mattiazzi Usaj, N. Sahin, H. Friesen, C. Pons, M. Usaj, M. P. D. Masinas, E. Shuteriqi, A. Shkurin, P. Aloy, Q. Morris, C. Boone, B. J. Andrews, Systematic genetics and single-cell imaging reveal widespread morphological pleiotropy and cell-to-cell variability. Mol. Syst. Biol. 16, e9243 (2020).

68. S. W. Michnick, E. D. Levy, The modular cell gets connected. Science 375, 1093–1094 (2022).

69. C. B. Brachmann, A. Davies, G. J. Cost, E. Caputo, J. Li, P. Hieter, J. D. Boeke, Designer deletion strains derived from Saccharomyces cerevisiae S288C: a useful set of strains and plasmids for PCR-mediated gene disruption and other applications. Yeast 14, 115–132 (1998).

70. H. E. Klock, S. A. Lesley, The Polymerase Incomplete Primer Extension (PIPE) method applied to high-throughput cloning and site-directed mutagenesis. Methods Mol. Biol. 498, 91–103 (2009).

71. D. K. Breslow, D. M. Cameron, S. R. Collins, M. Schuldiner, J. Stewart-Ornstein, H. W. Newman, S. Braun, H. D. Madhani, N. J. Krogan, J. S. Weissman, A comprehensive strategy enabling high-resolution functional analysis of the yeast genome. Nat. Methods 5, 711–718 (2008).

72. O. Ronneberger, P. Fischer, T. Brox, U-Net: Convolutional Networks for Biomedical Image Segmentation. doi: 10.48550/arXiv.1505.04597 (2015).

73. A. X. Lu, T. Zarin, I. S. Hsu, A. M. Moses, YeastSpotter: accurate and parameter-free web segmentation for microscopy images of yeast cells. Bioinformatics 35, 4525–4527 (2019).

74. O. Matalon, A. Steinberg, E. Sass, J. Hausser, E. D. Levy, Reprogramming protein abundance fluctuations in single cells by degradation. *bioRxiv*, doi: 10.1101/260695 (2018).

75. Maynes, S., F. Yang, and A. M. Parkhurst, Hysteresis: Tools for Modeling Rate-Dependent Hysteretic Processes and Ellipses (2013).

76. L. Komsta, Processing data for outliers. The Newsletter of the R Project Volume 6/2, May 2006 6, 10 (2006).

77. A. Fitzgibbon, M. Pilu, R. B. Fisher, Direct least square fitting of ellipses. [Preprint] (1999). 10.1109/34.765658.

78. J. Schindelin, I. Arganda-Carreras, E. Frise, V. Kaynig, M. Longair, T. Pietzsch, S. Preibisch, C. Rueden, S. Saalfeld, B. Schmid, J.-Y. Tinevez, D. J. White, V. Hartenstein, K. Eliceiri, P. Tomancak, A. Cardona, Fiji: an open-source platform for biological-image analysis. Nat. Methods 9, 676–682 (2012).

79. J.-Y. Tinevez, N. Perry, J. Schindelin, G. M. Hoopes, G. D. Reynolds, E. Laplantine, S. Y. Bednarek, S. L. Shorte, K. W. Eliceiri, TrackMate: An open and extensible platform for single-particle tracking. Methods 115, 80–90 (2017).

80. S. Malitsky, C. Ziv, S. Rosenwasser, S. Zheng, D. Schatz, Z. Porat, S. Ben-Dor, A. Aharoni, A. Vardi, Viral infection of the marine alga Emiliania huxleyi triggers lipidome remodeling and induces the production of highly saturated triacylglycerol. New Phytol. 210, 88–96 (2016).

81. J. Zeng, X. Huang, L. Zhou, Y. Tan, C. Hu, X. Wang, J. Niu, H. Wang, X. Lin, P. Yin, Metabolomics Identifies Biomarker Pattern for Early Diagnosis of Hepatocellular Carcinoma: from Diethylnitrosamine Treated Rats to Patients. Sci. Rep. 5, 16101 (2015).

82. J. Willforss, A. Chawade, F. Levander, NormalyzerDE: Online Tool for Improved Normalization of Omics Expression Data and High-Sensitivity Differential Expression Analysis. J. Proteome Res. 18, 732–740 (2019).

83. J. Josse, F. Husson, missMDA: A Package for Handling Missing Values in Multivariate Data Analysis. J. Stat. Softw. 70, 1–31 (2016).

84. R. Kolde, Pheatmap: pretty heatmaps. https://github.com/raivokolde/pheatmap.

85. J. Jumper, R. Evans, A. Pritzel, T. Green, M. Figurnov, O. Ronneberger, K. Tunyasuvunakool, R. Bates, A. Žídek, A. Potapenko, A. Bridgland, C. Meyer, S. A. A. Kohl, A. J. Ballard, A. Cowie, B. Romera-Paredes, S. Nikolov, R. Jain, J. Adler, T. Back, S. Petersen, D. Reiman, E. Clancy, M. Zielinski, M. Steinegger, M. Pacholska, T. Berghammer, S. Bodenstein, D. Silver, O. Vinyals, A. W. Senior, K. Kavukcuoglu, P. Kohli, D. Hassabis, Highly accurate protein structure prediction with AlphaFold. Nature 596, 583–589 (2021).

86. M. Mirdita, K. Schütze, Y. Moriwaki, L. Heo, S. Ovchinnikov, M. Steinegger, ColabFold: making protein folding accessible to all. Nat. Methods 19, 679–682 (2022).

87. W. M. Babinchak, B. K. Dumm, S. Venus, S. Boyko, A. A. Putnam, E. Jankowsky, W. K. Surewicz, Small molecules as potent biphasic modulators of protein liquid-liquid phase separation. Nat. Commun. 11, 5574 (2020).

88. S-Trap^TM^, *ProtiFi*. https://protifi.com/pages/s-trap.

89. J. Cox, M. Mann, MaxQuant enables high peptide identification rates, individualized p.p.b.-range mass accuracies and proteome-wide protein quantification. Nat. Biotechnol. 26, 1367–1372 (2008).

90. UniProt Consortium, UniProt: the universal protein knowledgebase in 2021. Nucleic Acids Res. 49, D480–D489 (2021).

91. J. E. Elias, S. P. Gygi, Target-decoy search strategy for increased confidence in large-scale protein identifications by mass spectrometry. [Preprint] (2007). 10.1038/nmeth1019.

92. Q. G. Gianetto, S. Wieczorek, Y. Couté, T. Burger, A peptide-level multiple imputation strategy accounting for the different natures of missing values in proteomics data, bioRxiv (2020). 10.1101/2020.05.29.122770.

93. M. E. Ritchie, B. Phipson, D. Wu, Y. Hu, C. W. Law, W. Shi, G. K. Smyth, limma powers differential expression analyses for RNA-sequencing and microarray studies. [Preprint] (2015). 10.1093/nar/gkv007.

94. E. D. Wong, S. R. Miyasato, S. Aleksander, K. Karra, R. S. Nash, M. S. Skrzypek, S. Weng, S. R. Engel, J. M. Cherry, Saccharomyces genome database update: server architecture, pan-genome nomenclature, and external resources. Genetics 224 (2023).

